# Peptides from non-immune proteins target infections through antimicrobial and immunomodulatory properties

**DOI:** 10.1101/2024.03.25.586636

**Authors:** Marcelo D. T. Torres, Angela Cesaro, Cesar de la Fuente-Nunez

## Abstract

Encrypted peptides have been recently described as a new class of antimicrobial molecules. They have been proposed to play a role in host immunity and as alternatives to conventional antibiotics. Intriguingly, many of these peptides are found embedded in proteins unrelated to the immune system, suggesting that immunological responses may extend beyond traditional host immunity proteins. To test this idea, here we synthesized and tested representative peptides derived from non-immune proteins for their ability to exert antimicrobial and immunomodulatory properties. Our experiments revealed that most of the tested peptides from non-immune proteins, derived from structural proteins as well as proteins from the nervous and visual systems, displayed potent *in vitro* antimicrobial activity. These molecules killed bacterial pathogens by targeting their membrane, and those originating from the same region of the body exhibited synergistic effects when combined. Beyond their antimicrobial properties, nearly 90% of the peptides tested exhibited immunomodulatory effects, modulating inflammatory mediators such as IL-6, TNF-α, and MCP-1. Moreover, eight of the peptides identified, collagenin 3 and 4, zipperin-1 and 2, and immunosin-2, 3, 12, and 13, displayed anti-infective efficacy in two different preclinical mouse models, reducing bacterial infections by up to four orders of magnitude. Altogether, our results support the hypothesis that peptides from non-immune proteins may play a role in host immunity. These results potentially expand our notion of the immune system to include previously unrecognized proteins and peptides that may be activated upon infection to confer protection to the host.

## Introduction

The innate immune system traditionally has been thought to be orchestrated by immune proteins. Computational and AI tools have recently been used to identify novel antimicrobial molecules encoded within genomes and proteomes^1–9;^ the presence of these molecules suggests that there is much dark matter still to be explored for antibiotic discovery. Encrypted peptides (EPs) were discovered as a result of such efforts. These molecules have now been identified across the tree of life, including in the human proteome^8,10,11^. Many EPs have been found to have potent antimicrobial properties^13^. Intriguingly, many of these peptides are derived from non-immune proteins. For instance, many human proteins with defined physiological roles, such as thrombin, protein C3, lactoferrin, pepsinogen A, signal peptide, CUB and EGF-like domain-containing protein, von Willebrand factor, and apolipoproteins, were demonstrated to harbor precursor forms of antimicrobial peptides that are released through cleavage by either host or bacterial proteases^8,10,12,14–17^. Altogether, these findings led us to formulate the “Meta Hypothesis” to test the notion that immunological response may be orchestrated by proteins and peptides beyond classical immune proteins.

Here, to test this hypothesis and assess whether encrypted peptides serve as a link between host immunity and other systems in the body, we explored and characterized representative peptides from the human proteome. Candidate sequences, totaling 39, were derived from both non-immune (26) and immune (13) related proteins. They were then ranked by our scoring function^8^ and according to their physicochemical features, including net charge, hydrophobicity, and length, which are known to influence antibacterial activity. The highest ranked candidates were screened against bacterial pathogens *in vitro* and in animal models. We analyzed their physicochemical features, elucidated their structure and mechanism of action, and evaluated their synergistic potential. Our findings revealed that most of the peptides tested displayed antimicrobial and immunomodulatory properties, thus providing a link between non-immune proteins and host immunity and supporting the “Meta Hypothesis”. These results suggest that non-immune proteins, whose functions are seemingly unrelated to host defense, may harbor peptides that contribute to immunological responses to invading pathogens, which are becoming increasingly prevalent^18,19^.

## Results

### Selection and synthesis of encrypted peptides from non-immune proteins

In our previous work mining the human proteome^8^, we found that just 1.7% of encrypted peptides were derived from immune proteins, whereas 98.3% came from non-immune proteins (**Fig. 1A**)^20^. EPs from the human proteome were previously classified^8^ as: (1) peptides derived from secreted proteins with diverse biological functions, including plasma proteins, coagulation factors, protein inhibitors, enzymes, and signaling cascade factors; or (2) peptide hormones with well-defined putative functions, but previously undisclosed antibiotic properties, such as neuropeptides, regulators of G-protein-coupled receptors (GPCRs), and diuretic hormones.

**Figure 1.**
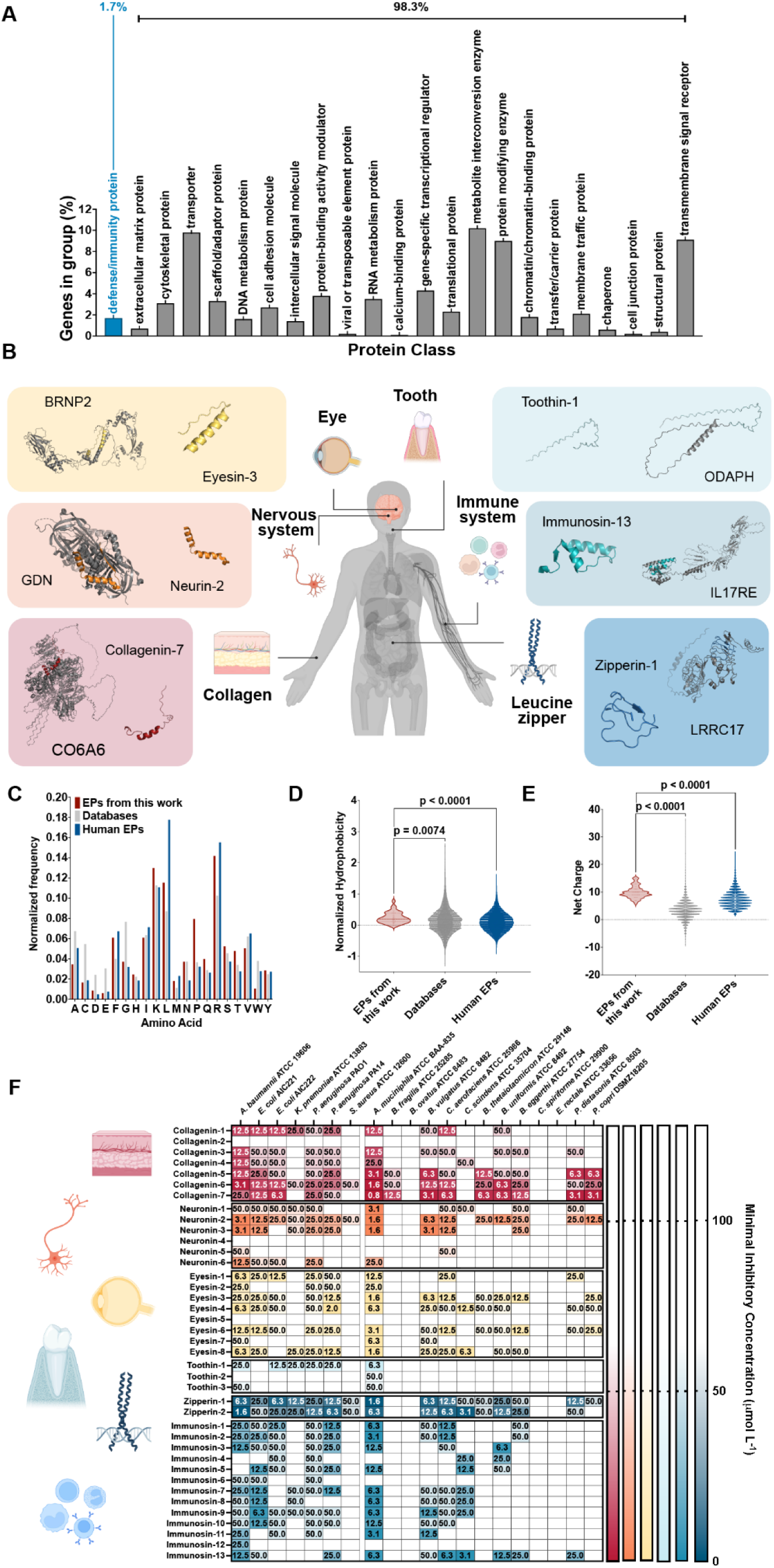
Origin, physicochemical properties, and antimicrobial activity of encrypted peptides. **(A)** Normalized abundance of genes encoding proteins containing predicted encrypted peptides. The analysis was performed using Panther Proteins Classification system with a false discovery rate cutoff of 0.05. **(B)** Tertiary structure of the host proteins and the selected encrypted peptides from 6 biogeographical systems: collagen- (red), nervous system- (orange), eye- (yellow), tooth- (light blue), immune system- (blue), and leucine zipper- (dark blue) related proteins. **(C)** Comparison of the amino acid frequency in the EPs from this work, with human EPs and well-known AMPs from databases. The amino acid counts were assessed using EPs from the human proteome, peptides from DBAASP, APD, and DRAMP (see also **Fig. S1A-B**). The distribution is shown for two of the most important physicochemical features for their antimicrobial activity: **(D)** normalized hydrophobicity and **(E)** net charge of EPs from this work, EPs from the human proteome and AMPs from databases (see also **Fig. S1C-H**). Hydrophobicity directly influences the interactions of the peptide with lipids from the bacterial membrane, while net charge directly influences the initial electrostatic interactions between the peptide and negatively charged bacterial membranes**. (F)** Minimal inhibitory concentration (MIC) values were obtained by testing peptide concentrations ranging from 0.78 to 50 μmol L^−1^ against 20 bacterial strains, i.e., seven pathogens and 13 human gut commensals. Reported data refer to assays carried out in triplicate and the heatmap shows the mode of MIC values collected from the experimental replicates. Figure created with BioRender.com.

In this study, we explored six groups of proteins from different biogeographical areas of the human body, which were classified as either 1) derived from non-immune proteins; or 2) from immune proteins **(Table S1)**. Immune proteins were included as controls. These included immunoglobulin superfamily member 10 (IGSF10), pro-interleukin-16 (IL16), leukocyte cell-derived chemotaxin-2 (LECT2), C-C motif chemokine 26 (CCL26), T-cell surface glycoprotein CD8 beta chain (CD8B), C-C motif chemokine 20 (CCL20), and interleukin-17 receptor E (IL17R). In our study, we included the following non-immune proteins: (1) collagen, one of the most abundant classes of proteins in humans and one that can be found in epithelial, connective, muscular, bone, and other structural tissues, e.g., collagen alpha-1(XXIV) chain (COOA1) involved in the processes of extracellular matrix organization and collagens biosynthesis; (2) nervous system-related proteins, ranging from large protein receptors to short peptide hormones, e.g., brain-derived neurotrophic factor (BDNF), which is an important signaling molecule, promoting the survival and differentiation of neuronal populations cells; (3) proteins from the eyes, known for their special relationship with the immune system since the eye is immune privileged^21^ [e.g. BMP/retinoic acid-inducible neural-specific protein 3 (BRNP3), whose overexpression seems to promote pituitary gonadotrope cell proliferation and migration], (4) proteins from the teeth, which are mostly structural proteins, e.g., odontogenesis associated phosphoprotein (ODAPH), which promotes the nucleation of hydroxyapatite in teeth; (5) leucine zipper proteins, DNA-binding and regulatory proteins with a leucine zipper domain that is made up of two motifs – a basic region that recognizes a specific DNA sequence and a series of leucine residues spaced seven residues apart along an α-helix (leucine zipper) that mediate dimerization [e.g. leucine-rich repeat-containing protein 17 (LRRC17), involved in regulating the bone homeostasis].

These protein groups were selected based on their high ranking, as determined by our scoring function^8,10,15,22^. Subsequently, representative EPs with physicochemical features of interest that are relevant for antimicrobial activity^23^ were chosen based on previous knowledge of EPs from the human proteome and potential for chemical synthesis through solid-phase peptide synthesis. Highly hydrophobic peptides and those containing multiple cysteine residues were filtered out from the selection process. In total, 39 EPs were selected for synthesis and biological characterization and named according to their parent protein groups: Immune proteeins (i.e, immunosins) and non-immune proteins [i.e., (1) collagenins, (2) neuronins, (3) eyesins, (4) toothins, and (5) zipperins].

### Composition and physicochemical features of encrypted peptides

The EPs selected for synthesis and biological characterization are representative sequences, meaning that they share a composition similar to those of other sequences from the human proteome^6,8^ (**Fig. 1B**). The main compositional differences between the peptides selected here compared to other human EPs^8^ are their higher frequency of proline (**Fig. 1B**), as well as overall higher content of aromatic residues (**Fig. S1A**) and lower content of aliphatic residues (**Fig. S1A**), making them less hydrophobic (**Fig. S1B**). When compared to known antimicrobial peptides (AMPs) available in databases such as the Database of Antimicrobial Activity and Structure of Peptides (DBAASP)^24^, the Antimicrobial Peptides Database (APD3)^25^, and the Data Repository of Antimicrobial Peptides (DRAMP)^26^, the EPs studied here exhibited a higher content of polar uncharged and basic residues and a lower content of acidic residues. In addition to their composition, we analyzed some of the most relevant physicochemical features that were used or directly affected by the scoring function ranking process. To extract the physicochemical features of the EP sequences (**Figs. 1C,D, S1C-H**), we used the properties tool in the DBAASP server^24^.

The selected EPs presented a higher mean net positive charge compared to AMPs and previously described EPs (**Fig. 1C**), and on average, were as hydrophobic as known AMPs, which, in turn, are more hydrophobic than previously described human EPs (**Fig. 1D**). The compositional similarities to AMPs and higher net positive charge account for the high hit rate obtained and the broad-spectrum activity profile of the selected EPs from all six biogeographical regions. Hydrophobicity, net charge values, and the amino acid composition of the EPs also influence other physicochemical features that are relevant for their biological functions. EPs have amphiphilicity (**Fig. S1C**) and propensity to *in vitro* aggregation (**Fig. S1D**) very similar to those of AMPs but different from previously described EPs. On average, the EPs from the six biogeographical regions exhibited a higher normalized hydrophobic moment (**Fig. S1E**), but lower isoelectric point (**Fig. S1F**), angle subtended by the hydrophobic residues (**Fig. S1G**), and disordered conformation propensity (**Fig. S1H**) than AMPs and other human EPs.

### Antimicrobial activity assays

To assess the potential antimicrobial activity of the selected peptides from the human proteome, we treated clinically relevant pathogens (*Acinetobacter baumannii*, *Escherichia coli*, *Klebsiella pneumoniae*, *Pseudomonas aeruginosa*, and *Staphylococcus aureus*) and species representative of the most abundant commensal bacteria from the gut microbiome [*Akkermansia muciniphila*, *Bacteroides eggerthi*, *Bacteroides fragilis*, *Bacteroides thetaiotaomicron*, *Bacteroides uniformis*, *Bacteroides vulgatus* (*Phocaeicola vulgatus*), *Collinsella aerofaciens*, *Clostridium scindens*, *Eubacterium rectale*, *Parabacteroides distasonis*, and *Prevotella copri*] to EPs at a gradient of concentrations ranging from 0.78 to 50 μmol L^-1^. Thirty-six of the total 39 selected EPs (92.3%) were active against at least one of the bacterial strains tested at the range of concentrations analyzed (0.78 to 50 μmol L^-1^; **Fig. 1E**). These data underscored the accuracy of our selection process based on physicochemical features and amino acid composition. The three inactive peptides were derived from proteins from three different biogeographical areas: collagenin-2, neuronin-4, and eyesin-5. As observed for previously reported EPs^8^, most of the EPs showed broad-spectrum antimicrobial activity, regardless of the biogeographical region of origin. Thirty-five of 39 (89.7%) were active against pathogens and 34 of 39 (87.2%) were active against commensals. The most active groups were collagenins, immunosins, and zipperins. Interestingly, toothin-2 was only active against the gut commensal *A. muciniphila* (at 50 μmol L^-1^). While two immune system-derived EPs were selective toward pathogens, immunin-12 was selective to *A. baumannii* (at 25 μmol L^-1^) and immunin-6 was selective to the pathogens *A. baumannii*, *E. coli* AIC221 and *P. aeruginosa* PAO1 (all at 50 μmol L^-1^). Our results confirmed that EPs, unlike other natural peptides with known antimicrobial activity^27^, can effectively kill not only pathogens but also target commensal strains. It is noteworthy that the commensals *B. ovatus*, *C. spiriforme*, and *E. rectale* were not susceptible to any of the EPs tested.

Taken together, these findings strengthen our hypothesis that encrypted fragments of proteins with diverse functions can be part of the immune response and may reflect an evolutionary mechanism that expands protein capabilities while minimizing genomic expansion.

### Secondary structure studies

Because of their amphiphilicity, net charge, hydrophobicity, and length, peptides with antimicrobial activity mostly assume an α-helical conformation upon contact with bacterial membranes. Considering that the selected EPs have different amino acid residue compositions and physicochemical features compared to standard AMPs, we sought to investigate their secondary structure. To determine the secondary structure tendencies of the EPs from different biogeographical regions, we analyzed them in a helix-inducing medium, trifluoroethanol in water (3:2, v:v), to mimic their interaction with the bacterial membrane (**Figs. 2A-B, S2A-F**). Most of the EPs tested were predominantly unstructured in solution (**Fig. S2**); the only family with all unstructured peptides was the toothins (**Fig. S2D**). Leucine zipper proteins present many well characterized helical domains, and as expected, the zipperins showed high helical content (**Figs. 2A-B, S2F**). Collagenins-1 and −6, neuronin-2, and eyesin-5 were the other EPs that were mostly helical out of the non-immune system-related proteins (**Fig. 2A**). Immunosin-10 was the peptide from the immune system with the highest helical content (62%), whereas immunosins 4 and 11 were the other two peptides from immune system-related proteins that presented relatively high helical content (50 and 51%, respectively). Interestingly, immunosin-12 exhibited a predominant β-antiparallel structure (100%), which is common for constrained AMPs but much less frequent than α-helical conformations (**Fig. 2A**). We attribute this structure to the high number of proline residues and the long amphipathic sequence of immunosin-12: RPLLLLAYFSRLCAKGDIPPPLRALPRYRLLRDLPRLLR.

**Figure 2.**
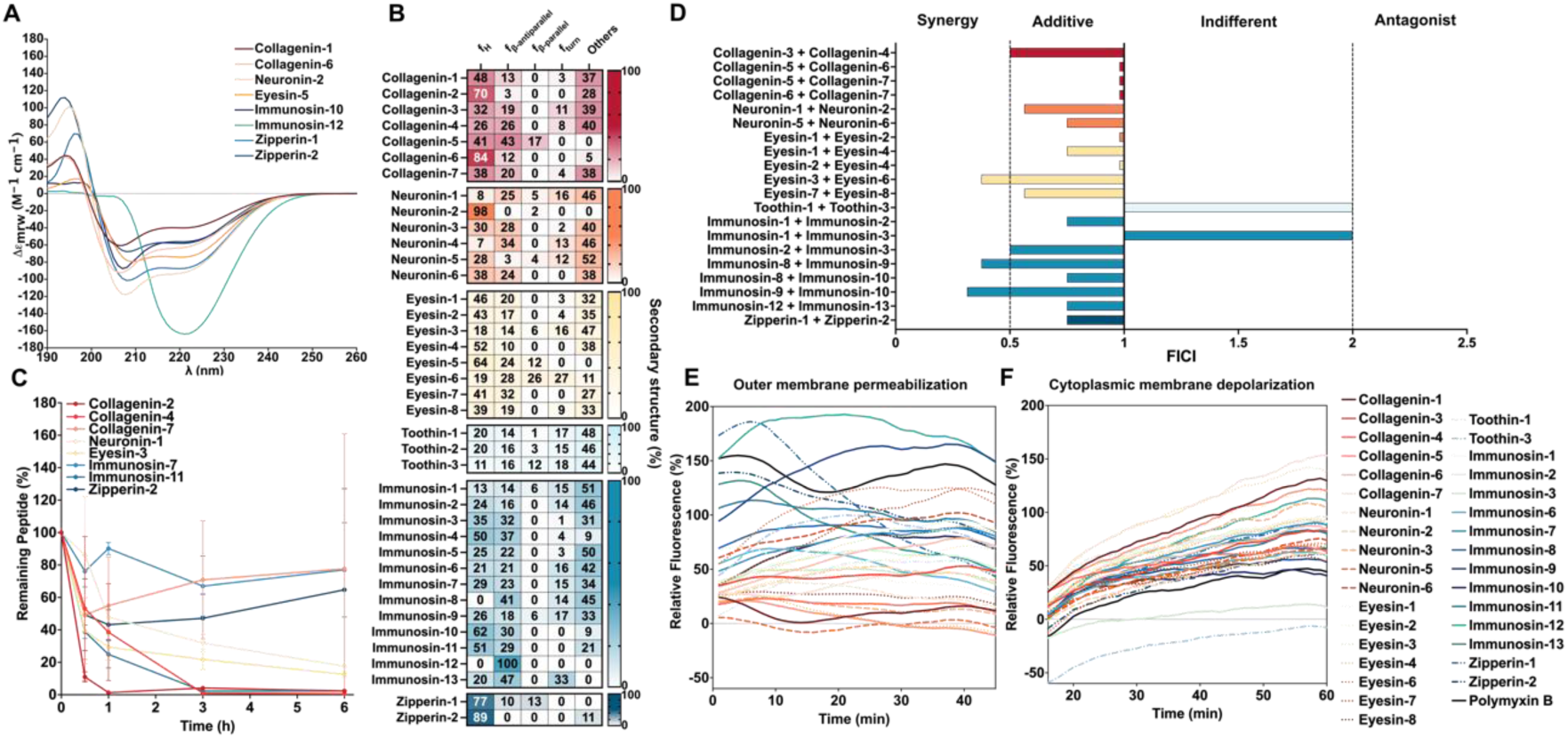
Secondary structure, resistance to proteolytic degradation, synergy, and mechanism of action of encrypted peptides. **(A)** Circular dichroism spectra of the most active EPs from various biogeographical systems and **(B)** percentage of secondary structure in trifluoroethanol and water mixture (3:2, v/v)^44,45^ (see also **Fig. S2**). Unlike most of the active AMPs^23,30,32^, EP activity is not directly correlated with α-helical content. **(C)** The resistance to proteolytic degradation of representative EPs from each biogeographical system was assessed by exposing the peptides for six hours to human serum. Aliquots of the resulting solution were analyzed by liquid chromatography coupled to mass spectrometry; the most active peptides were those with higher resistance to proteolytic degradation. **(D)** The synergistic interaction between pairs of EPs from the same biogeographical system was analyzed by checkerboard assays with 2-fold serial dilutions starting at 2×MIC to MIC/32. The histogram shows the fractional inhibitory concentration index (FICI) values obtained for each pair of Eps. Low FICI values (≤0.5) represent synergistic interactions, intermediary values (0.5<FICI≤1) and higher values represent indifferent (1<FICI≤2) and antagonistic (<2) interactions (see also **Fig. S3**). To assess whether EPs act on the bacterial membrane, all active EPs against *A. baumannii* ATCC 19606 were tested in outer membrane permeabilization and cytoplasmic membrane depolarization assays. The fluorescent probe 1-(N-phenylamino)naphthalene (NPN) was used to indicate membrane permeabilization **(E)** caused by the EPs tested (see also **Fig. S4**). The fluorescent probe 3,3′-dipropylthiadicarbocyanine iodide (DiSC_3_-5) was used to indicate membrane depolarization **(F)** caused by collagenins, neuronins, eyesins, toothins, immunosins, and zipperins (see also **Fig. S5**). The values shown represent the relative fluorescence of both probes used as a non-linear fitting using as baseline the untreated control (buffer + bacteria + fluorescent dye) in comparison to polymyxin B, an antibiotic known for its permeabilization properties that also shows some depolarizing activity. All experiments were performed in three independent replicates.

### Resistance to proteolytic degradation assays

Next, we selected representatives peptides derived from immune and non-immune proteins with different activity profiles and the most different sequence composition to study their resistance to proteolytic degradation in human serum. The most active EPs from collagen and immune system proteins resisted degradation for 6 h with up to 77% of their initial concentration remaining (**Fig. 2C**). Up to 65% of the initial concentration of zipperin-2, one of the most active peptides tested, remained after 6 h of continuous exposure to human serum proteases (**Fig. 2C**). Collagenin-2, the only inactive EP tested, degraded rapidly: after 30 min only 11% remained (**Fig. 2C**).

### Synergy assays

To investigate whether EPs from non-immune and immune proteins can synergize and thus potentiate each other’s activity against pathogens, we performed checkerboard assays^8,10^ at peptide concentrations ranging from twice the MIC to concentrations up to 64-fold lower in the same conditions as used for the antimicrobial assays. First, we selected the EPs that belong to proteins that co-exist and were active against *A. baumannii* ATCC 19606 (**Fig. 1B**), an opportunistic nosocomial pathogen with increasing antibiotic resistance that has resulted in significant mortality worldwide^28^. Most of the combinations tested resulted in synergistic or additive interactions, calculated by using the fractional inhibitory concentration index^29^ (FICI, **Fig. 2D**) leading to concentrations in the same order of magnitude as the ones observed for the expression levels of the parent proteins (**Fig. S3**). Five of the pairs synergized, i.e., FICI ≤ 0.5; for example, (1) eyesins 3 and 6 from the protein BMP/retinoic acid-inducible neural-specific protein 2, which is involved in the cell cycle regulation of neuronal cell (FICI = 0.375), and (2) the two pairs of immunosins: 8 and 9 (FICI = 0.375) and 9 and 10 (FICI = 0.3125) from the protein T-cell surface glycoprotein CD8 beta chain, which is involved in the immunoregulatory interaction in responses against both external and internal signals. Collagenins 3 and 4 from collagen alpha-1(XVI) chain, involved in signal transduction and extracellular matrix organization, and immunosins 2 and 3 from protein immunoglobulin superfamily member 10, which has a role in the control of the early migration of neurons, also synergized (FICI = 0.5), though not as much as the abovementioned pairs. The two pairs that showed indifferent interactions were toothins 1 and 3 from the odontogenesis-associated phosphoprotein and immunosins 1 and 3 from immunoglobulin superfamily member 10, with FICI values of 2.

### Mechanism of action studies

To investigate whether EPs from the six biogeographical regions operate by targeting the bacterial membrane, we performed fluorescence assays. We selected all antimicrobial hits (**Fig. 1B**) against the pathogens and assessed the ability of EPs (at their MIC value) to permeabilize the outer membrane (**Figs. 2E**, **S4A-F**) and/or depolarize the cytoplasmic membrane (**Figs. 2F, S5A-F**) of bacterial cells. First, to assess whether EPs permeabilized the outer membrane of *A. baumannii*, we performed 1-(N-phenylamino)naphthalene (NPN) uptake assays. NPN is a lipophilic dye whose fluorescence increases in hydrophobic environments, but it is not able to permeate the bacterial outer membrane unless the membrane is damaged. As a positive control we used the antibiotic polymyxin B (PMB), a highly active cyclic peptide known to permeate Gram-negative outer membranes^8^. Collagenins showed lower permeabilization of the outer membrane compared to PMB (**Figs. 2E, S4A**), followed by neuronins (**Figs. 2E, S4B**), of which the only peptide with relatively high permeabilization was neuronin-6. Toothins (**Figs. 2E, S4D**) and eyesins (**Figs. 2E, S4C**) showed intermediary permeabilization compared to PMB. Eyesins 6 and 7 permeabilized membranes as effectively as PMB. The strongest permeabilizers were the non-immune derived compounds zipperins 1 and 2 (**Figs. 2E, S4F**) and the immune protein derived molecules immunosins 9 and 12 (**Figs. 2E, S4E**), which presented low mean hydrophobicity (0.09-0.15) and an intermediary net positive charge (7-9), except for immunosin-9 (0.77 and 15, respectively). In summary, the compounds derived from immune and non-immune proteins assessed in this study permeabilized the outer membrane of bacteria to levels similar to previously reported EPs^6,8,10^ and AMPs ^30,31^.

Next, we used 3,3′-dipropylthiadicarbocyanine iodide (DiSC_3_-5), a fluorophore that is utilized to determine depolarization of the cytoplasmic membrane of bacteria reflected by changes in the membrane transmembrane potential. DiSC_3_-5 is first accommodated in the membrane lipid bilayer and then migrates to the extracellular environment, generating fluorescence. Interestingly, the non-immune protein derived collagenins (**Figs. 2F, S5A**) that did not permeabilize the outer membrane showed the highest depolarizing properties followed by neuronins 1 and 3 (**Figs. 2F, S5B**), toothin-1 (**Figs. 2F, S5D**), eyesins 1, 3, and 4 (**Figs. 2F, S5C**), and zipperin-1 (**Figs. 2F, S5F**). Immunosins 1 and 12 (**Figs. 2F, S5A**) also caused very low depolarization of the cytoplasmic membrane. Collagenins have low mean hydrophobicity (0.06-0.21) and net positive charge ranging from 8 to 11, which are common ranges for peptides with relatively long sequences (33-48 residues) and amphiphilic amino acid composition (**Figs. 1D-E, S1C-H**). Immunosin-12 was the only compound, considering both immune and non-immune proteins studied here, that effectively permeabilized (**Figs. 2E, 2F, S4E, S5E**) and depolarized the outer and cytoplasmic membrane.

Overall, these data suggest that while there is no clear correlation between the physicochemical properties of EPs analyzed in our study and their mechanism of action, we can observe certain tendencies among groups of EPs from different biogeographical regions: some EPs tend to operate predominantly by permeabilizing or depolarizing the outer and cytoplasmic membrane. These results are intriguing as they demonstrate that there is much more to EPs than what was previously known^8,10^. This finding sheds new light on the complexity and diversity of EPs, highlighting the need for further research in this area to fully understand their potential as antimicrobial agents.

### *In vitro* cytotoxicity and hemolytic activity assays

To evaluate their effects on eukaryotic cells, we tested whether the EPs would damage human red blood cells (RBCs), immortalized human keratinocytes (HaCaT), and PMA-differentiated human monocytes (THP-1). Overall, the EPs had higher cytotoxic effects when assayed on RBCs than on HaCaT or PMA-differentiated THP-1 cells **(Figs. 3B-D, S6)**. Eyesin-2, 3, and 6, derived from BMP/retinoic acid-inducible neural-specific proteins (BRNP2 and 3), and toothin-1, 2, and 3, all peptides derived from odontogenesis-associated phosphoprotein (ODAPH), displayed HC_50_ (hemolytic concentration causing 50% cell death) values equal or even lower than 10 µmol L^-1^. Contrarily, it was observed that all immune-related peptides exhibited values exceeding 25 µmol L^-1^ **(Figs. 3B, S6)**. Indeed, 31 of the 39 EPs (79.5%) tested with the RBCs (including all immunosins) showed MIC_100_ values lower than the respective HC_50_ values **(Fig. 3B)**. On the other hand, all EPs tested with HaCaT and PMA-differentiated THP-1 cells showed CC_50_ (cytotoxic concentration causing 50% cell death) values higher than 20 and 10 µmol L^-1^, respectively **(Figs. 3C-D, S6)**. Particularly, zipperin-1 and 2, derived from leucine-rich repeat-containing protein 17 (LRRC17), and immunosin-12, derived from interleukin-17 receptor E (IL17RE), did not affect the viability of keratinocytes even when tested at the highest concentration (50 µmol L^-1^). Thus, those three EPs were considered safe because their toxic concentration is at least 2 times higher than their antimicrobial activity **(Figs. 3C, S6)**. Overall, no significant differences in cytotoxic effects were observed between immune and non-immune derived peptides when tested on HaCaT or PMA-differentiated THP-1 cell lines.

**Figure 3.**
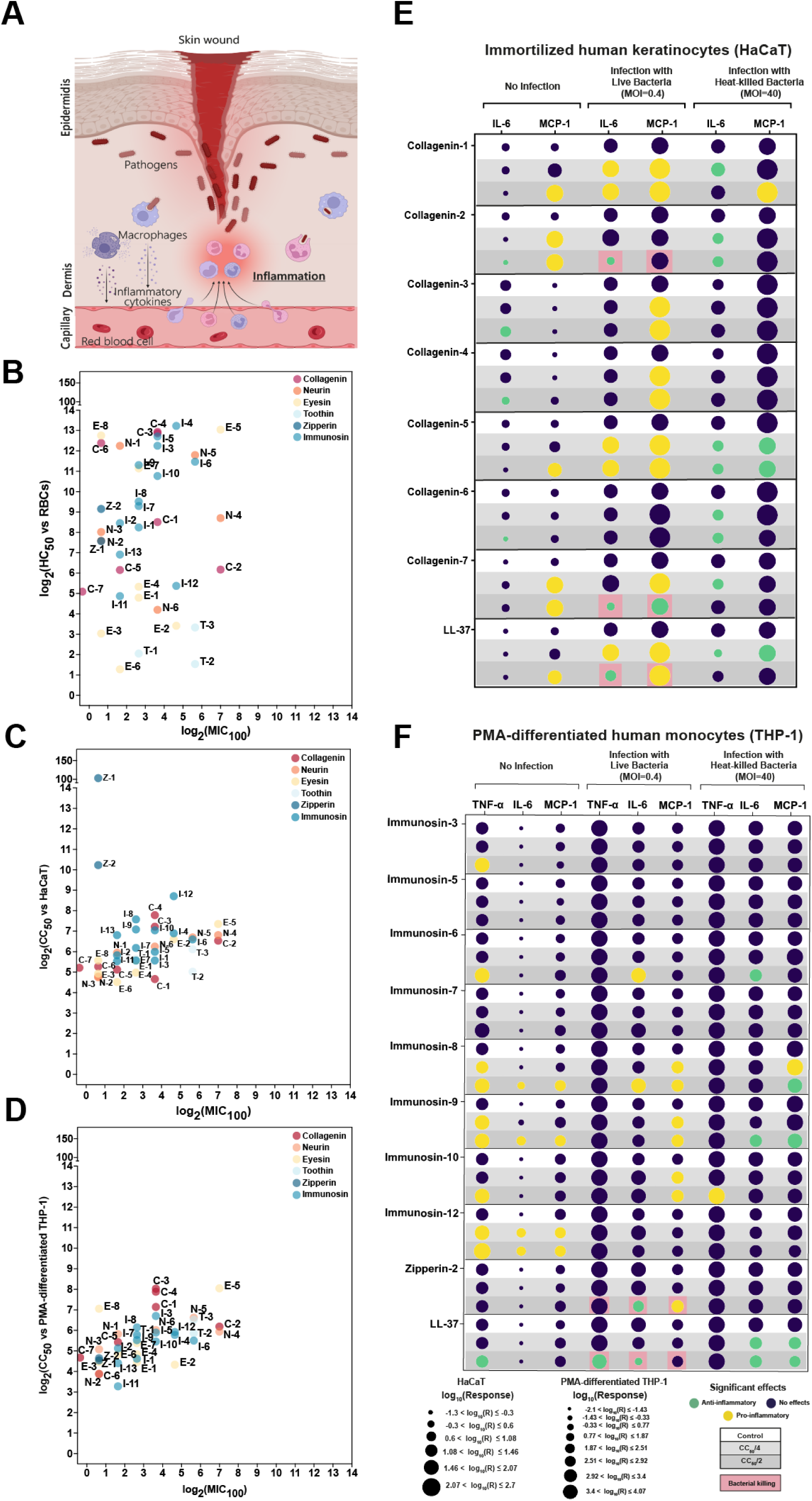
Cytotoxicity and immunomodulatory properties of encrypted peptides. **(A)** Schematic representation of skin wound infection triggering inflammatory response. Correlations between cytotoxicity and antimicrobial activity – expressed as MIC (minimal inhibitory concentration for complete bacterial killing) – are shown in scatter plots. The impact on **(B)** red blood cells (RBCs) in terms of hemolytic effects, and cytotoxicity for **(C)** immortalized human keratinocytes (HaCaT) and **(D)** PMA-differentiated human monocytes (THP-1) are expressed using predicted HC_50_ (hemolytic concentrations leading to 50% cell death) and CC_50_ values (cytotoxic concentrations leading to 50% cell death). HC_50_ and CC_50_ values were obtained through interpolation of the dose-response using a non-linear regression curve (see also **Fig. S6**). The release of pro-inflammatory markers from **(E)** HaCaT and **(F)** PMA-differentiated THP-1 cells, whether uninfected or infected with live or heat-killed *A. baumannii*, was examined after a 20 h treatment with each EP. Samples were collected and analyzed to detect levels of IL-6, TNF-α, and MCP-1. Data are shown as mean of two independent experiments (in quadruplicate) for each group (see also **Fig. S7-S10**). Figure created with BioRender.com.

### Immunomodulatory assays

To assess whether the identified peptides modulated the immune response, we tested their effects on cells uninfected or infected with live or heat-killed *A. baumannii* ATCC 19606. EPs identified in collagen-related proteins (collagenins 1-7) were tested on keratinocytes, while EPs derived from immune system- and leucine zipper-related proteins (namely immunosins 3, 5, 6, 7, 8, 9, 10, 12 and zipperin-2) were tested on PMA-differentiated THP-1. Almost all collagenins were found to increase the levels of the monocyte chemoattractant protein-1 (MCP-1) when tested on uninfected HaCaT or the same cells infected with live *A. baumannii* cells **(Figs. 3E, S7-S8)**. Interestingly, collagenin-2, which is derived from disintegrin and metalloproteinase domain-containing protein 12 (ADAM12), decreased the levels of IL-6 in the absence or presence of infections with both live and heat-killed bacteria **(Figs. 3E, S7)**. In contrast, collagenin-5, derived from collagen alpha-6(VI) chain (CO6A6), a protein involved in collagen biosynthesis, displayed pro-inflammatory effects in the absence of infection and in the presence of live bacteria and significantly reduced the levels of IL-6 and MCP-1 when the infection was caused by heat-killed bacteria. Thus, collagenin-5 appears to trigger diverse inflammatory responses under varying infection conditions. **(Figs. 3E, S8)**.

In contrast, immunosin-6, 8, 9, 10, and 12, derived from leukocyte cell-derived chemotaxin-2 (LECT2), C-C motif chemokine 26 (CCL26), T-cell surface glycoprotein CD8 beta chain (CD8B), and interleukin-17 receptor E (IL17RE), increased the level of TNF-α, IL-6 and MCP-1 when tested on PMA-differentiated THP-1 cells uninfected and stimulated with live bacteria **(Figs. 3F, S9-S10)**. Immunosin-6, derived from LECT2, and immunosin-8 and 9, derived from CD8B, reduced IL-6 and MCP-1 levels in cells infected with heat-killed bacteria, displaying an anti-inflammatory effect **(Fig. 3F, S9-S10)**. Zipperin-2, derived from leucine-rich repeat-containing protein 17 (LRRC17), decreased IL-6 while increasing MCP-1 levels. Those anti-inflammatory effects might be explained by the concomitant antimicrobial activity observed in the same experimental condition **(Figs. 3F, S10)**.

Overall, similar to the immune-related EPs, the non-immune derived EPs tested appear to play a role in both pro- and anti-inflammatory effects, depending on the specific physiological context. Interestingly, in over 70% of the cases, the tested EPs, such as collagenins, zipperins and immunosins, were found to increase the levels of MCP-1, thus indicating a significant function in immune cell recruiting.

### Anti-infective activity of encrypted peptides in preclinical animal models

To assess whether the selected EPs maintained their *in vitro* antimicrobial activity in complex living systems, we tested them in two different mouse models (**Fig. 4A**). Collagenins were tested in a skin abscess infection model^30,32,33^, and zipperins and immunosins were tested in a preclinical murine deep thigh infection model^6,8,34^.

**Figure 4.**
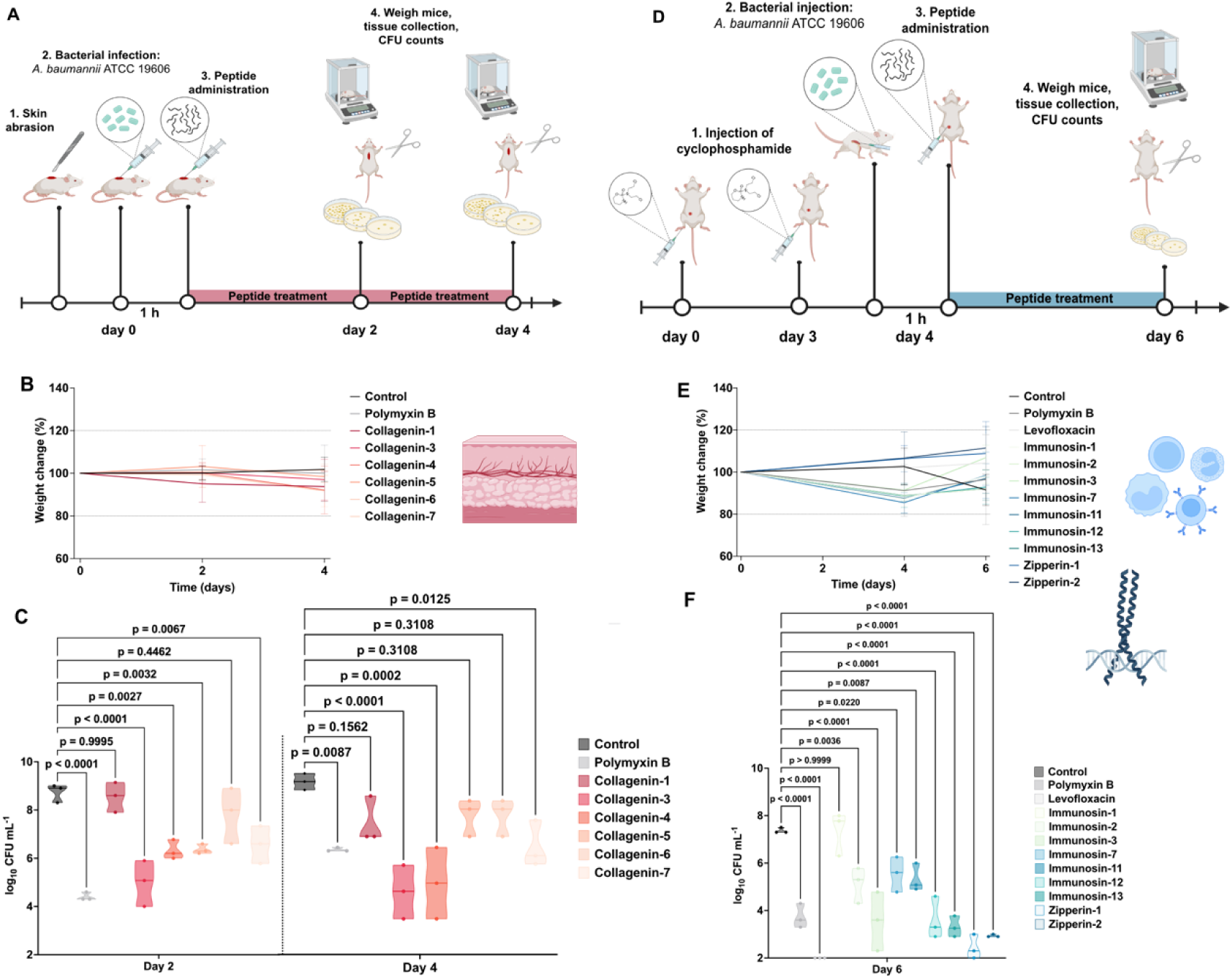
Anti-infective activity of encrypted peptides in preclinical animal models. **(A)** Schematic of the skin abscess infection mouse model used to assess the anti-infective activity of the collagenins against *A. baumannii* cells in wounded epidermis. In this model, mice had their back shaved, wounded, and infected with a load of *A. baumannii* cells. They were treated one hour after the infection was established with a single dose of the collagenins at their MIC for *A. baumannii* ATCC 19606. **(B)** Weight changes were used as a proxy for toxicity of the EPs during the whole extent of the experiment. We considered up to 20% weight change as the threshold for lack of toxicity. **(C)** Mice were euthanized two- and four-days post-infection. Each group (treated and untreated) consisted of three mice (n = 3) and the bacterial loads used for infection of each mouse came from different inocula grown and prepared independently. Collagenin-3 presented bactericidal effects comparable to those of the positive control antibiotic, polymyxin-B. Collagenins 4, 5, and 7 had bacteriostatic effect two days after infection. Four days post-infection, the only peptides that did not allow proliferation of *A. baumannii* cells were collagenins 3 and 4 (see also **Fig. S11**). **(D)** Schematic of the deep thigh infection mouse model used to assess the anti-infective activity of immunosins and zipperins against *A. baumannii* cells in an intramuscular infection. In this model, mice were first rendered neutropenic by two rounds of treatment of cyclophosphamide. Subsequently, bacterial cells were injected intramuscularly in the right thigh followed by the intraperitoneal administration of a single dose of one of the two families of EPs at their MIC against *A. baumannii*. Two days post-infection (six days after the start of the experiment), mice were euthanized. Each group had three subjects (n = 3), with independently prepared bacterial loads. **(E)** During the entire experiment, weight change was monitored. **(F)** Except for immunosin-1, all peptides tested were active. Immunosins 2, 7, and 11 showed bacteriostatic activity, while immunosins 3, 12, and 13 and zipperins 1 and 2 had bactericidal activity comparable to that observed for polymyxin-B. Levofloxacin, another antibiotic used as positive control, was the most active compound tested, leading to clearance or significant reduction of the bacterial load. One-way ANOVA followed by Dunnett’s test was used to determine statistical significance in panels **(C)** and **(F)**, and the respective p-values are presented for each group. All groups were compared to the untreated control, and the violin plots display the median and upper and lower quartiles. Data in **(B)** and **(E)** are the mean plus and minus the standard deviation. Figure created with BioRender.com.

For the skin abscess infection experiments, we selected all collagenins with *in vitro* activity (**Fig. 1B**) against *A. baumannii* cells and little or no toxicity (**Fig. 3A**) for eukaryotic cells (collagenins 1, 3, 4, 5, 6, and 7). *A. baumannii* (2×10^6^ cells in 20 μL) was administered to a skin abscess created on the shaved back of each mouse (n=3, **Fig. 4A**). One dose of PMB (control) or collagenins was administered to the infected area at their MIC. All the peptides except collagenins 1 and 6 showed bactericidal activity two days post-infection. Collagenin-3 (MIC = 12.5 μmol L^-1^) had bactericidal effects as high as those of PMB in the skin abscess model (**Fig. 4B**). Four days post-infection, collagenins 3 and 4 were still as active as PMB (**Fig. 4B**). The activity levels of the active collagenins were comparable to those of some of the most potent AMPs described to date and tested with the same model, *i.e.*, polybia-CP^32^ and PaDBS1R6^35^. Mouse weight changes were used as a proxy for toxicity, no weight changes (i.e., no deleterious effects) were observed in the animals (**Fig. 4C**). Although no antimicrobial activity was detected *in vitro* for collagenin-2, derived from disintegrin and metalloproteinase domain-containing protein 12, this EP sterilized the wound infection when applied on the back of the mouse (**Fig. S11A-B**). The anti-infective activity of this EP seems to increase when it is in complex biological matrices, such as in the presence of cells or embedded in tissue. This indicates that the antimicrobial and immunomodulatory properties of collagenin-2 might be triggered by external biological factors.

For the deep thigh infection model (n = 3, **Fig. 4D**), we selected zipperins (1 and 2) and immunosins (1, 2, 3, 7, 11, 12, and 13) as they displayed *in vitro* activity against *A. baumannii* (**Fig. 1B**) at concentrations that were not toxic for eukaryotic cells (**Fig. 3D**). Bacterial loads of 5×10^6^ cells in 100 μL were injected intramuscularly, and one dose of the selected peptides was injected intraperitoneally at its MIC. Two-days post-treatment (6 days after the beginning of the experiment), the two peptides derived from non-immune proteins (zipperins 1 and 2) presented high bactericidal activity, similar to some of the immunosins (3, 12, and 13) and at levels comparable to PMB, but were not as active as levofloxacin (**Fig. 4E**). Immunosins 2, 7, and 11 showed bacteriostatic effects in these experiments. No significant changes in mouse weight were observed (**Fig. 4F**).

## Discussion

In this study, we have tested the hypothesis that non-immune proteins may contribute to host immunity by releasing peptide fragments possessing antimicrobial and immunomodulatory properties. Our results revealed that 92.3% of the peptides tested displayed activity against at least one bacterial strain.

Comparative analysis of physicochemical features revealed that the selected peptides shared similar traits with other human EPs; thus, physicochemical attributes like net charge and amphiphilicity significantly contributed to antimicrobial activity. The killing mechanisms of these compounds involved both membrane permeabilization and depolarization, with specific peptides demonstrating a preference for one of these mechanisms. For example, collagenins operated predominantly by depolarizing the cytoplasmic membrane. Furthermore, the combination of EPs derived from the same parental proteins, particularly eye proteins and immune-related proteins, demonstrated enhanced antimicrobial activity, i.e., synergistic antimicrobial interactions. Notably, most of the active encrypted peptides were not toxic for eukaryotic cells at their MIC. Interestingly, non-immune protein derived peptides (collagenins and zipperins) were found to modulate the release of inflammatory mediators, demonstrating an ability to modulate the immune response in addition to their direct antimicrobial activity.

The anti-infective activity of the most active and non-toxic peptides was validated in two different pre-clinical animal models. Collagenins-2, 3, and 4 significantly reduced the bacterial load in infected skin wounds, while zipperins-1 and 2 and immunosins-3, 12, and 13, when administered intraperitoneally, decreased intramuscular thigh infections. Overall, zipperin peptides from the leucine-rich repeat-containing protein 17 (LRRC17), mainly involved in bone homeostasis^36–38^, showed the highest anti-infective properties both *in vitro* and in an animal model. Immune proteins-related peptides derived from T-cell surface glycoprotein CD8 beta chain (CD8B)^39,40^, a protein already known to have a role in the immune response, exhibited antimicrobial and immunomodulatory properties. Interestingly, collagenin-2, a bioactive fragment identified in disintegrin and metalloproteinase domain-containing protein 12 (ADAM12), completely cleared the infection in the skin scarification model (**Fig. S11**), even though its *in vitro* antimicrobial activity was poor (**Fig. 1F**). However, this peptide reduced IL-6 levels and stimulated MCP-1 release (**Figs. 1E and S7**). These data align with recent studies that point to the role of ADAM12 in promoting immune cell filtration^41–43^.

Our results provide evidence in support of the “Meta Hypothesis”, whereby non-immune proteins may harbor peptide sequences with immune functions. Our results underscore the multifaceted nature of human proteins, many of which may play previously unrecognized roles in host immunity.

**STAR Methods:**

- Key resource table
- Resource availability

`o` Lead contact
`o` Materials availability
`o` Data and code availability
- Experimental model and subject details

`o` Bacterial strains and growth conditions
`o` Eukaryotic cells culture
`o` Red blood cells and human serum
`o` Skin abscess infection mouse model
`o` Deep thigh infection mouse model
- Method details

`o` Human proteome screening for encrypted peptides
`o` Peptide synthesis
`o` Minimal inhibitory concentration assays
`o` Circular dichroism experiments
`o` Resistance to proteolytic degradation assays
`o` Outer membrane permeabilization assays
`o` Cytoplasmic membrane depolarization assays
`o` Synergy assays
`o` Cytotoxicity assays
`o` Hemolysis assays
`o` Cell stimulation and detection of inflammatory markers by Enzyme-Linked Immunosorbent Assay (ELISA)
- Quantification and statistical analysis

`o` Reproducibility of the experimental assays
`o` Statistical tests
`o` Statistical analysis

## Acknowledgments

Cesar de la Fuente-Nunez holds a Presidential Professorship at the University of Pennsylvania and acknowledges funding from the Procter & Gamble Company, United Therapeutics, a BBRF Young Investigator Grant, the Nemirovsky Prize, Penn Health-Tech Accelerator Award, and the Dean’s Innovation Fund from the Perelman School of Medicine at the University of Pennsylvania. Research reported in this publication was supported by the Langer Prize (AIChE Foundation), the National Institute of General Medical Sciences of the National Institutes of Health under award number R35GM138201, and the Defense Threat Reduction Agency (DTRA; HDTRA11810041, HDTRA1-21-1-0014, and HDTRA1-23-1-0001). We thank Dr. Mark Goulian for kindly donating the following strains: *Escherichia coli* AIC221 [*Escherichia coli* MG1655 phnE_2::FRT (control strain for AIC222)] and *Escherichia coli* AIC222 [*Escherichia coli* MG1655 pmrA53 phnE_2::FRT (polymyxin resistant)]. We thank Dr. Gui-Shuang Ying from the Center for Preventive Ophthalmology and Biostatistics (CPOB) Consulting Service for advice on our statistical analyses. We thank Dr. Karen Pepper for editing the manuscript and de la Fuente Lab members for insightful discussions. Figures created with BioRender.com are attributed as such. Molecules were rendered using the PyMOL Molecular Graphics System, Version 2.5.2 Schrödinger, LLC.

## Author contributions

Conceptualization: MDTT, CFN

Methodology: MDTT, AC, CFN

Investigation: MDTT, AC

Visualization: MDTT, AC

Funding acquisition: CFN

Supervision: CFN

Formal analysis: MDTT, AC

Writing – original draft: MDTT, AC, CFN

Writing – review & editing: MDTT, AC, CFN

## STAR Methods

### Key resources table

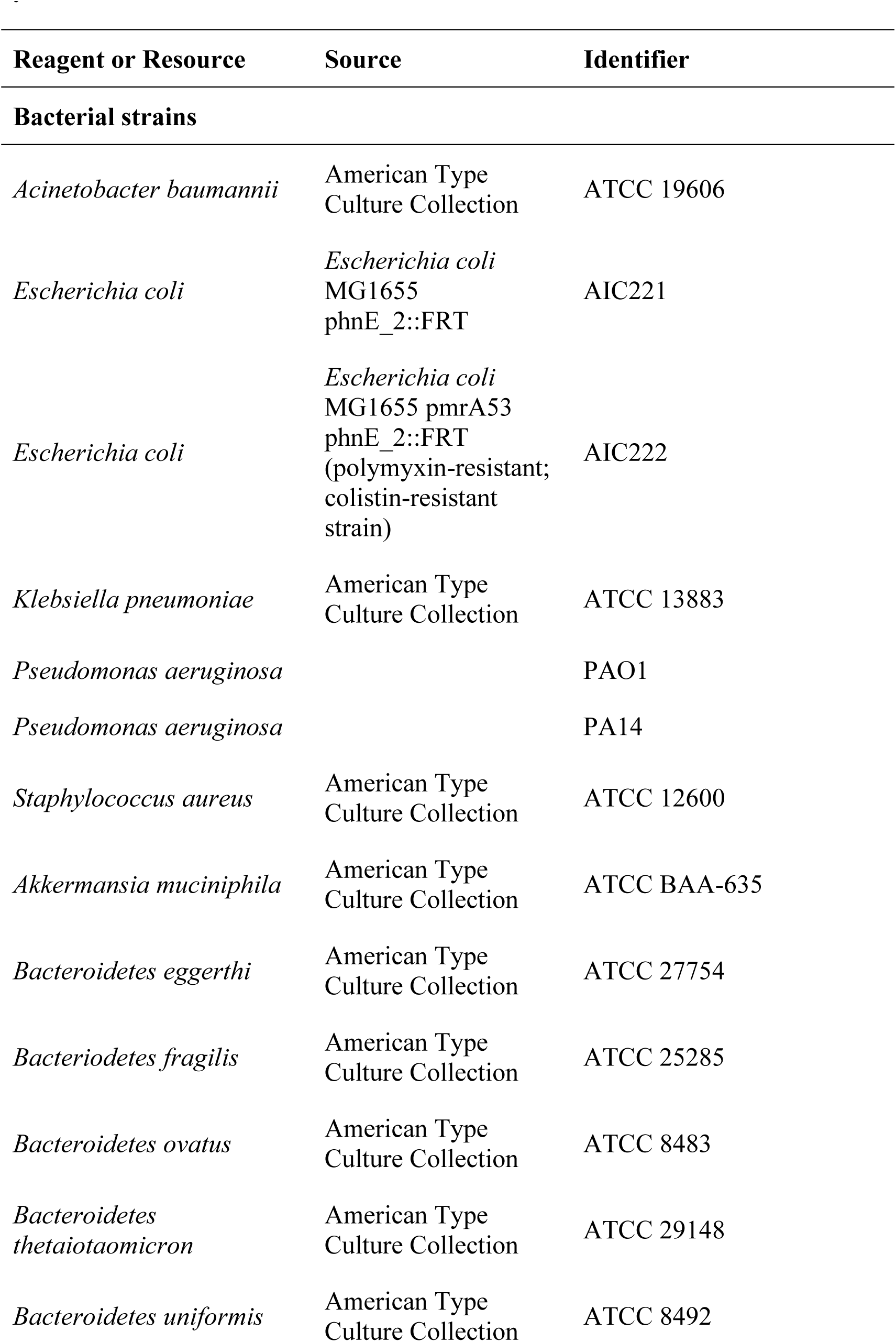
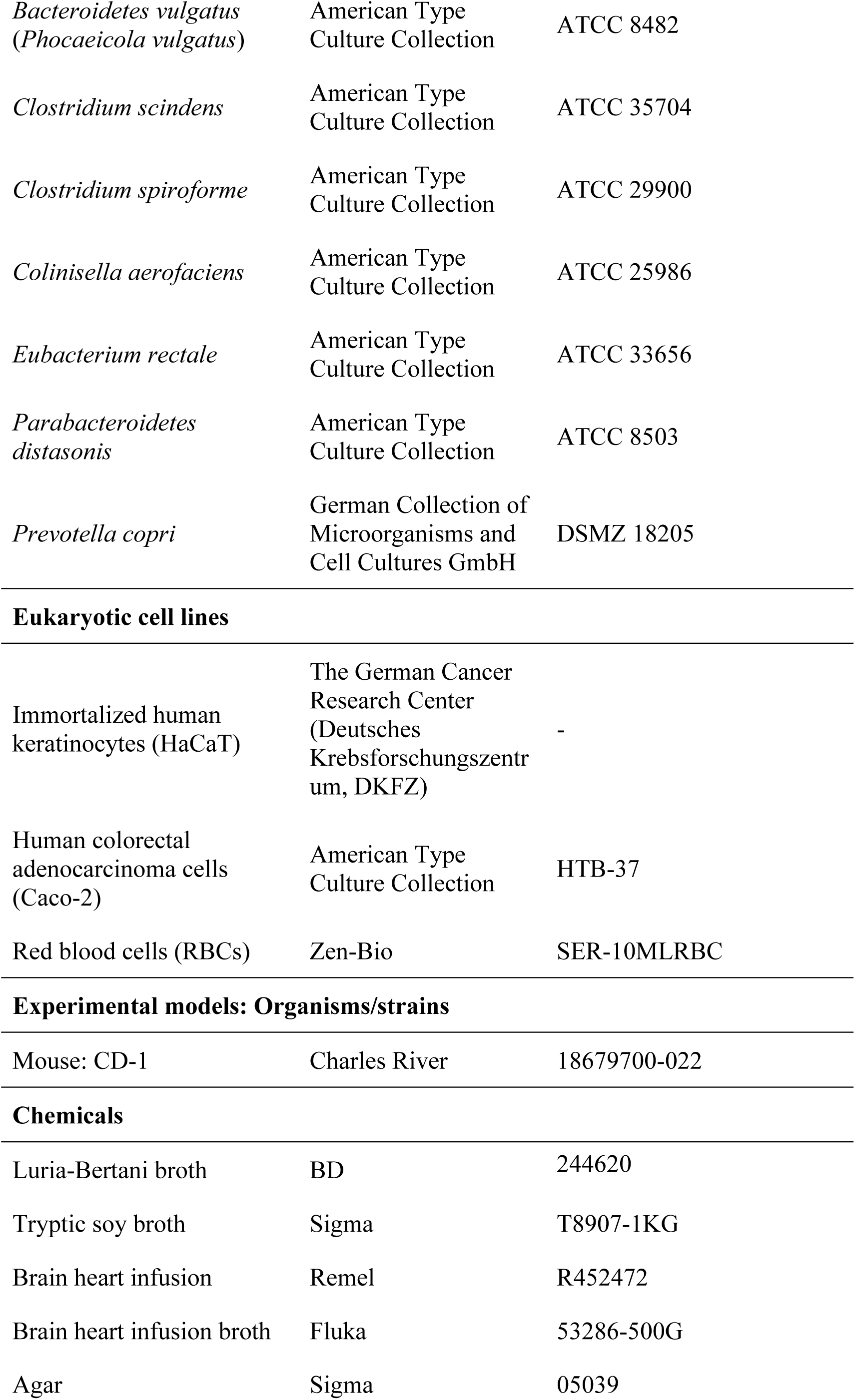
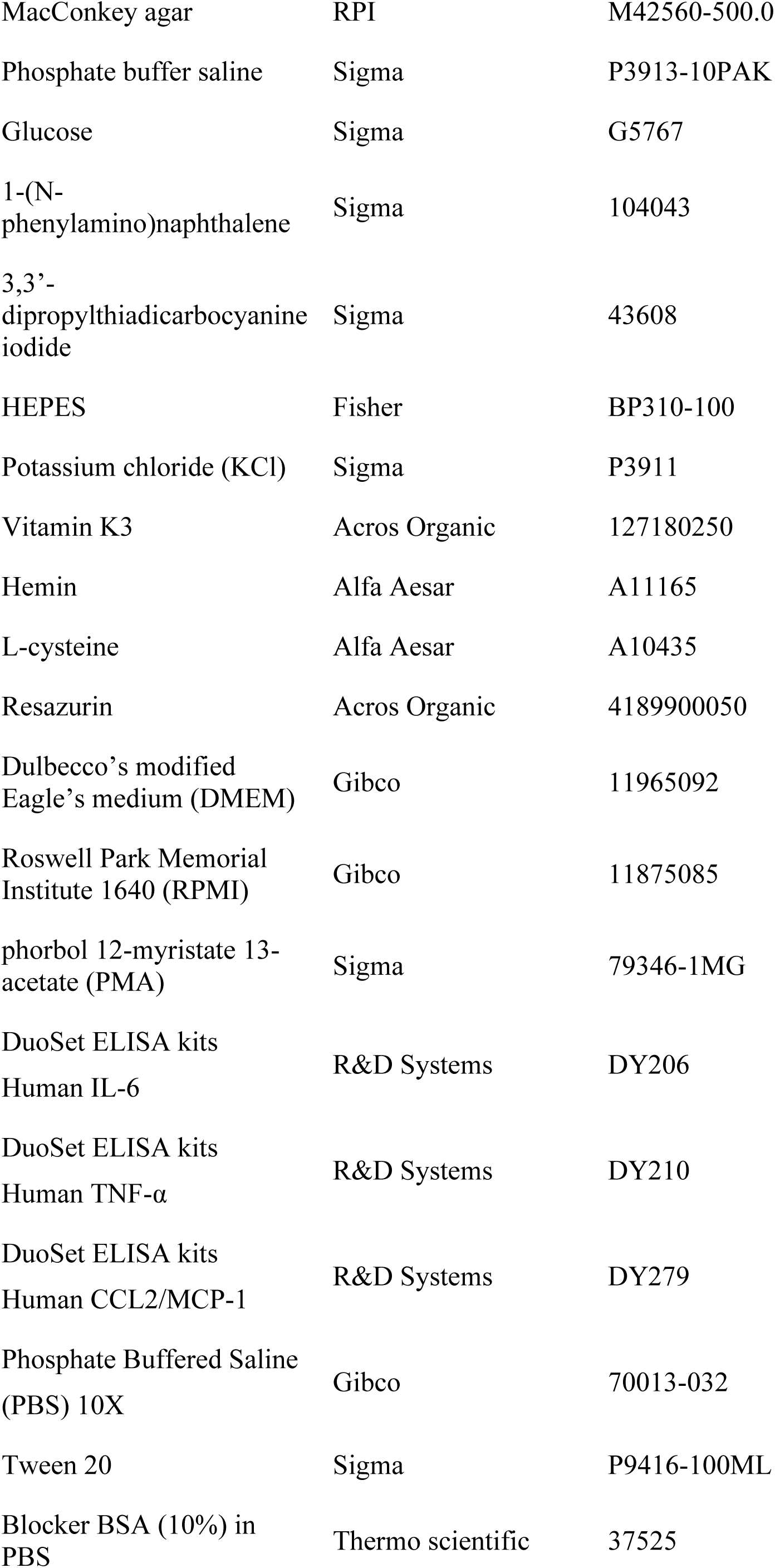
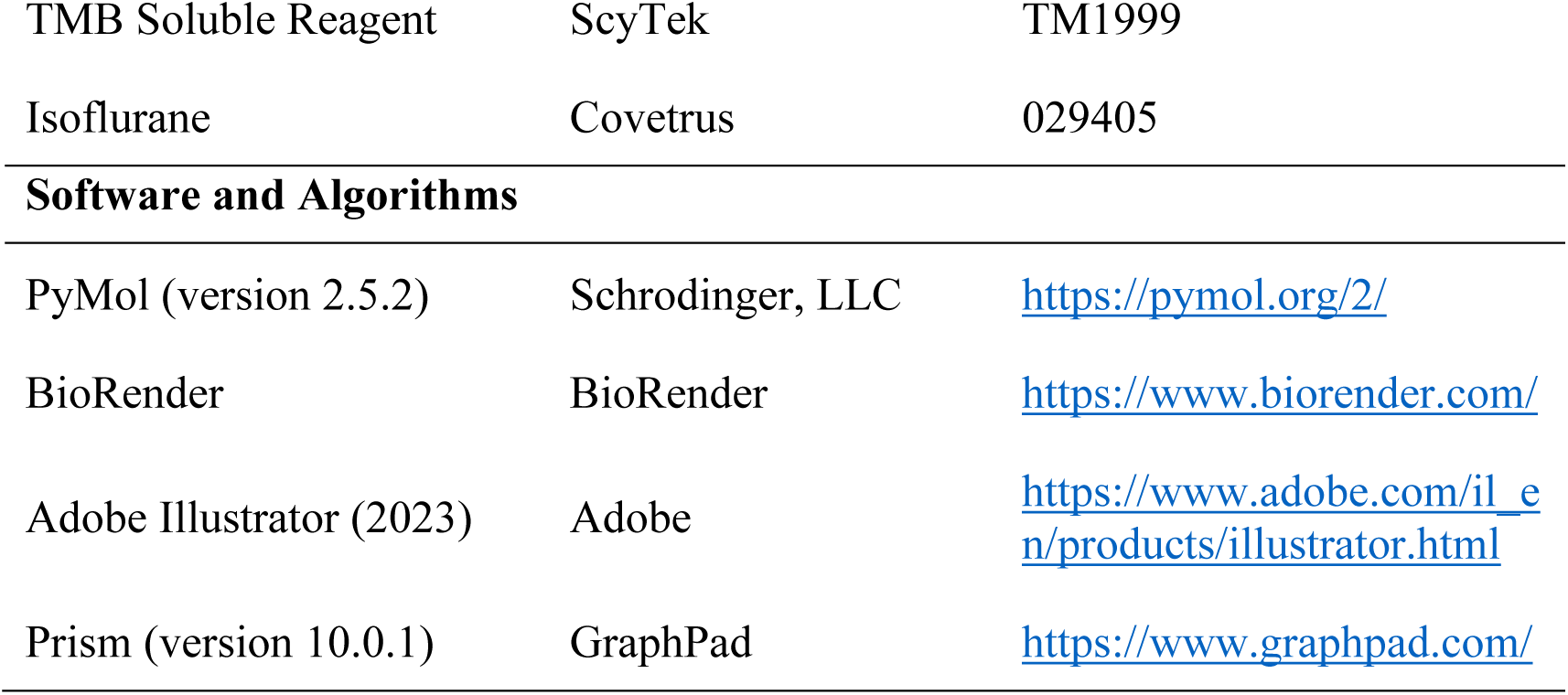

### Resource Availability

#### Lead contact

Further information and requests for resources should be directed to and will be fulfilled upon reasonable request by the lead contact, Cesar de la Fuente-Nunez.

#### Materials availability

This study did not generate new unique reagents.

#### Data and code availability

Test results used for analysis will be provided upon reasonable request.

- The human proteome data are publicly available at https://www.uniprot.org/.
- All original code has been deposited at GitHub in the link: https://gitlab.com/machine-biology-group-public/amp_scoring.
- Any additional information required to reanalyze the data reported in this paper is available from the lead contact upon request.

### Experimental model and subject details

#### Bacterial strains and growth conditions

In this study, we used the following pathogenic bacterial strains: *Acinetobacter baumannii* ATCC 19606, *Escherichia coli* AIC221 [*Escherichia coli* MG1655 phnE_2::FRT (control strain for AIC 222)] and *Escherichia coli* AIC222 [*Escherichia coli* MG1655 pmrA53 phnE_2::FRT (polymyxin resistant; colistin-resistant strain)], *Klebsiella pneumoniae* ATCC 13883, *Pseudomonas aeruginosa* PAO1, *Pseudomonas aeruginosa* PA14, and *Staphylococcus aureus* ATCC 12600. We also used commensal bacterial strains including: *Akkermansia muciniphila* ATCC BAA-635, *Bacteroides eggerthi* ATCC 27754, *Bacteroides fragilis* ATCC 25285, *Bacteroides thetaiotaomicron* ATCC 29148, *Bacteroides uniformis* ATCC 8492, *Bacteroides vulgatus* (*Phocaeicola vulgatus*) ATCC 8482, *Collinsella aerofaciens* ATCC 25986, *Clostridium scindens* ATCC 35704, *Eubacterium rectale* ATCC 33656, *Parabacteroidetes distasonis* ATCC 8503, and *Prevotella copri* DSMZ18205. All pathogens were grown in Luria-Bertani (LB) broth and on LB agar while the gut commensals were cultured using brain–heart infusion (BHI) broth and on BHI agar, both supplemented with 0.1% (v:v) vitamin K3 (1 mg mL^-1^), 1% (v:v) hemin (1 mg mL^-1^, diluted with 10 mL of 1 N sodium hydroxide), and 10% (v:v) _L_-cysteine (0.05 mg mL^-1^). Resazurin (250 mg L^-1^) was used as oxygen indicator. Pseudomonas Isolation (*Pseudomonas aeruginosa* strains) agar plates were exclusively used in the case of *Pseudomonas* species. In all the experiments, bacteria were inoculated from one-isolated colony and grown overnight (16 h) in liquid medium at 37 °C. In the following day, inoculums were diluted 1:100 in fresh media and incubated at 37 °C to mid-logarithmic phase^46,47^.

#### Eukaryotic cells culture

Immortalized human keratinocytes (HaCaT obtained from Deutsches Krebsforschungszentrum - DKFZ)^48^ and human monocytes (THP-1 ATCC TIB-202) were cultured in high-glucose Dulbecco’s modified Eagle’s medium (DMEM) and Roswell Park Memorial Institute 1640 (RPMI), respectively, previously supplemented with 1% antibiotics (penicillin/streptomycin) and 10% fetal bovine serum (FBS). Both cell lines were grown at 37 °C in a humidified atmosphere containing 5% CO_2_.

#### Red blood cells and human serum

Red blood cells (RBCs) and human serum were purchased separately from Zen-Bio. The samples were obtained from the same certified healthy donor (blood type A^-^).

#### Skin abscess infection mouse model

The back of six-week-old female CD-1 mice under anesthesia were shaved and injured with a superficial linear skin abrasion made with a needle. An aliquot of *A. baumannii*

ATCC 19606 (2×10^6^ CFU mL^-1^; 20 μL) previously grown in LB medium until 0.5 OD mL^-1^ (optical value at 600 nm) and then washed twice with sterile PBS (pH 7.4, 13,000 rpm for 3 min) was added to the scratched area. Peptides diluted in sterile water at MIC value were administered to the wound area 1 h after the infection. Two- and four-days post-infection animals were euthanized, and the scarified skin was excised, homogenized using a bead beater (25 Hz for 20 min), 10-fold serially diluted, and plated on McConkey agar plates for CFU quantification. The experiments were performed using three mice per group. The skin abscess infection mouse model was revised and approved by the University Laboratory Animal Resources (ULAR) from the University of Pennsylvania (Protocol 806763).

#### Deep thigh infection mouse model

Six-week-old female CD-1 mice were rendered neutropenic by two doses of cyclophosphamide (150 mg Kg^-1^ and 100 mg Kg^-1^) applied intraperitoneally 3 and 1 days before the infection. At day 4 of the experiment, the mice were infected in their right thigh by a 100 μL intramuscular injection of the *A. baumannii* ATCC19606 in PBS at concentration of 5×10^6^ CFU mL^-1^. The bacterial cells were grown in LB broth, washed twice with PBS solution, and diluted at the desired concentration. The peptides were administrated intraperitoneally two hours after the infection. Two-days post-infection mice were euthanized, and the tissue from the right thigh was excised, homogenized using a bead beater (25 Hz for 20 min), 10-fold serially diluted, and plated on McConkey agar plates for bacterial colonies counting. The experiments were performed using three mice per group. The deep thigh infection mouse model was revised and approved by the University Laboratory Animal Resources (ULAR) from the University of Pennsylvania (Protocol 807055).

### Method details

#### Human proteome screening for encrypted peptides

All canonical and isoform sequences of *Homo sapiens* proteins were downloaded from UniProt^49^. Our previously described Python script^8^ was updated to account for encrypted peptides containing the last amino acid residue of the proteins (https://gitlab.com/machine-biology-group-public/amp_scoring). Briefly, we used the updated script to scan all proteins using multiple moving windows with lengths ranging from 8 to 50 residues. Each sequence was scored using sequence length, net charge, and hydrophobicity that correlates with their propensity to present antimicrobial activity^8^. We filtered out identical peptide sequences and listed the top 1,000 scoring peptides per sequence length, yielding a 43,000-sequences dataset of candidates from across the whole human genome. An analogous search was performed to generate a final list of top candidates, but focusing solely on secreted proteins, as those were predicted to provide more biologically relevant encrypted peptides. After this process, we selected the highest ranked peptides from the six biogeographical regions of interest. Before proceeding to the solid-phase peptide synthesis, we filtered out sequences that were too hydrophobic or present hydrophobic clusters aiming to avoid problems such as aggregation, multiple recouplings, and low yields during the synthesis process.

#### Peptide Synthesis

All peptides used in the experiments were purchased from AAPPTec and synthesized by solid-phase peptide synthesis using the Fmoc strategy.

#### Minimal inhibitory concentration assays

Broth microdilution assays were performed to determine the minimum inhibitory concentration (MIC) values of each peptide. Peptides were added to nontreated polystyrene microtiter 96-well plates and 2-fold serially diluted in sterile water from 0 to 128 μmol L^-1^. Bacterial inoculum at 4×10^6^ CFU mL^-1^ in LB or BHI medium was mixed 1:1 with the peptide. The MIC was defined as the lowest concentration of peptide able to completely inhibit the bacterial growth after 24 h of incubation at 37 °C. All assays were done in three independent replicates.

#### Circular dichroism experiments

The circular dichroism experiments were conducted using a J1500 circular dichroism spectropolarimeter (Jasco) in the Biological Chemistry Resource Center (BCRC) at the University of Pennsylvania. Experiments were performed at 25 °C, the spectra graphed are an average of three accumulations obtained with a quartz cuvette with an optical path length of 1.0 mm, ranging from 260 to 190 nm at a rate of 50 nm min^-1^ and a bandwidth of 0.5 nm. The concentration of all EPs tested was 50 μmol L^-1^, and the measurements were performed in a mixture of water and trifluoroethanol (TFE) in a 3:2 ratio, with respective baselines recorded prior to measurement. A Fourier transform filter was applied to minimize background effects. Helical fraction values were calculated using the single spectra analysis tool on the server BeStSel^50^.

#### Resistance to proteolytic degradation assay

The resistance to enzymatic degradation was evaluated by incubating the encrypted peptides in 25% human serum in water. Briefly, peptides at a concentration of 2 mg mL^-1^ were exposed to an aqueous solution of 25% human serum obtained from Zen-Bio (male donor, blood type A-) for 6 h. Aliquots were collected after 0.5, 1, 3, and 6 h, and incubated for 10 minutes with 10 μL of trifluoroacetic acid. The samples were processed in a Waters Acquity ultra-performance liquid chromatography-mass spectrometry (UPLC-MS) equipped with a photodiode array detector (190-400 nm data collection) and a Waters TQD triple quadrupole MSMS, with 5 μL injections. The column used was a Waters Acquity UPLC HSS C_18_, 1.8 μm (2.1 mm x 50 mm). The mobile phases used were A (100% water with 0.1%, v/v, formic acid) and B (100% acetonitrile with 0.1%, v/v, formic acid), Fisher optima grades. Measurements were made by ionization ESI +/- simultaneous over m/z 100-2,000 Da. The percentage of remaining undamaged peptide was calculated by integrating the area under the curve related to the peptide at time point zero.

| Time (min) | A (%) | B (%) | Flow rate (mL min<sup>-1</sup>) |
| --- | --- | --- | --- |
| 0 | 95 | 5 | 0.5 |
| 0.5 | 95 | 5 | 0.5 |
| 2.5 | 5 | 95 | 0.5* |
| 3 | 5 | 95 | 0.5 |
| 3.25 | 5 | 95 | 0.5 |

\*linear gradient.

#### Outer membrane permeabilization assays

N-phenyl-1-napthylamine (NPN) uptake assay was used to evaluate the ability of the peptides to permeabilize the bacterial outer membrane. Inocula of *A. baumannii* ATCC 19606 were grown to an OD at 600 nm of 0.4 mL^-1^, centrifuged (10,000 rpm at 4 °C for 10 min), washed and resuspended in 5 mmol L^-1^ HEPES buffer (pH 7.4) containing 5 mmol L^-1^ glucose. The bacterial solution was added to a white 96-well plate (100 μL per well) together with 4 μL of NPN at 0.5 mmol L^-1^. Consequently, peptides diluted in water were added to each well, and the fluorescence was measured at λ_ex_ = 350 nm and λ_em_ = 420 nm over time for 45 min. The relative fluorescence was calculated using the untreated control (buffer + bacteria + fluorescent dye) and polymyxin B (positive control) as baselines and the following equation was applied to reflect % of difference between the baselines and the sample:

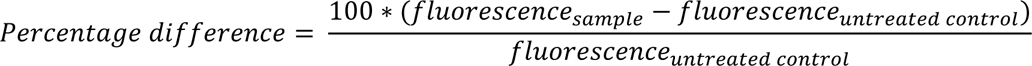

#### Cytoplasmic membrane depolarization assays

The cytoplasmic membrane depolarization assay was performed using the membrane potential-sensitive dye 3,3’-dipropylthiadicarbocyanine iodide (DiSC3-5). *A. baumannii* ATCC 19606 and *P. aeruginosa* PAO1 in the mid-logarithmic phase were washed and resuspended at 0.05 OD mL^-1^ (optical value at 600 nm) in HEPES buffer (pH 7.2) containing 20 mmol L^-1^ glucose and 0.1 mol L^-1^ KCl. DiSC3-5 at 20 μmol L^-1^ was added to the bacterial suspension (100 μL per well) for 15 min to stabilize the fluorescence which indicates the incorporation of the dye into the bacterial membrane, and then the peptides were mixed 1:1 with the bacteria to a final concentration corresponding to their MIC100 values. Membrane depolarization was then followed by reading changes in the fluorescence (λ_ex_ = 622 nm, λ_em_ = 670 nm) over time for 60 min. The relative fluorescence was calculated using the untreated control (buffer + bacteria + fluorescent dye) and polymyxin B (positive control) as baselines and the following equation was applied to reflect % of difference between the baselines and the sample:

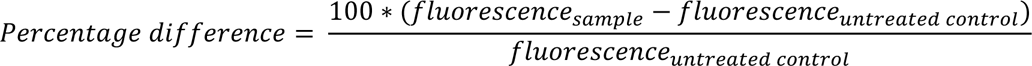

#### Synergy assays

Combinations of two encrypted peptides from the same protein were tested on *A. baumannii* ATCC 19606 strains using the checkboard assay. Briefly, two-fold serial dilutions of each peptide were orthogonally mixed and incubated with bacterial suspension at the final concentration of 2×10^6^ CFU mL^-1^ in LB for 24 h at 37 °C. The fractional inhibitory concentration indexes (FICI) were defined in order to attribute whether the interactions between peptides were synergistic (FICI≤0.5), additive (0.5>FICI≥1), or indifferent (FICI>1).

#### Cytotoxicity assays

HaCaT cells were seeded into 96-well plates at the density of 5×10^3^ cells per well one day before the treatment with increasing concentrations of peptide (3.12-50 μmol L^-1^). THP-1 monocytes were differentiated into macrophages by incubation with 100 nmol L-^1^ phorbol 12-myristate 13-acetate (PMA) for three days before the incubation with the peptides. After the incubation with each peptide for one day, we performed a MTT assay. Briefly, the MTT reagent was dissolved at 0.5 mg mL^-1^ in medium (DMEM and RPMI in the case of HaCaT and PMA-differentiated THP-1, respectively) without phenol red was used to replace cell culture supernatants (100 μL per well), and the samples were incubated for 4 h at 37 °C to obtain the insoluble formazan salts. The resulting salts were then solubilized in 0.04 mol L^-1^ hydrochloric acid (HCl) in anhydrous isopropanol and quantified by measuring the absorbance at 570 nm using a spectrophotometer.

#### Hemolysis assays

To evaluate the release of hemoglobin from human erythrocytes upon treatment of each of the encrypted peptides, human red blood cells (RBCs) were obtained from ZenBio (male donor, blood type A^-^) obtained from heparin anti-coagulated blood. RBCs were washed with PBS (pH 7.4) four times by centrifugation at 800 g for 10 min. Aliquots of 200-fold diluted cells (75 μL) were mixed with peptide solution (0.78-100 μmol L^-1^; 75 μL), and the mixture was incubated for 4 h at room temperature. After the incubation, the plate was centrifuged at 1,300 g for 10 min to precipitate cells and debris, and 100 μL of supernatant from each well were transferred to a new 96-well plate for absorbance reading (405 nm) using an automatic plate reader. The percentage of hemolysis was defined by comparison with negative control (samples containing PBS) and positive control [samples containing 1% (v/v) SDS in PBS solution].

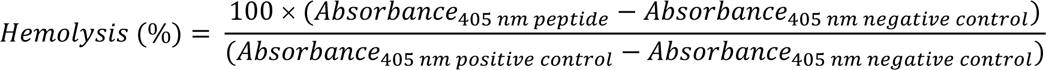

#### Cells stimulation and detection of inflammatory markers by Enzyme-Linked Immunosorbent Assay (ELISA)

HaCaT and PMA-differentiated THP-1 were plated at the density of 50,000 cells per well (100 μL) and washed twice with PBS before incubation with peptide alone or co-incubation with peptide and live or heat-killed *A. baumannii* ATCC 19606. A culture of *A. baumannii* at exponential growth phase was washed twice with PBS by centrifuging the samples at 5,000 g for 5 min. To inactivate the bacteria, the sample was exposed at 75 °C for 30 min. Live and heat-killed bacteria were then diluted in tissue culture medium at multiplicity of infection (MOI) 0.4 and 40, respectively. Following incubation with peptides and/or bacteria, cytokines levels (IL-6, TNF-α, and MCP-1) within the medium were quantify by using human immunoassay kits (DuoSet ELISA kits, R&D Systems, Minneapolis, MN) according to the manufacturer’s instructions. Optical density of each sample was measured by using an ELISA reader set to 450 nm with a wavelength correction set to 540 nm. To detect antimicrobial effects displayed by the peptides towards live bacteria cells, the amount of *A. baumannii* cells in each supernatant collected was quantified by measuring the absorbance at 600 nm and by plating the sample on agar plates.

### Quantification and statistical analysis

#### Reproducibility of the experimental assays

Unless otherwise stated, all assays were performed in three independent biological replicates as indicated in each figure legend and Experimental Models and Methods details sections. The values obtained for cytotoxic activity were estimated by non-linear regression based on the screen of peptides in a gradient of concentrations and represent the hemolytic and the cytotoxic concentration values needed to lyse and kill 50% of the cells present in the experiment. For the cytotoxic activity assays, two technical replicates were performed within each of the three biological replicates. The ELISA assays were performed in two biological replicates with four technical replicates per experiment. In the skin abscess and thigh infection mouse models, we used three mice per group following established protocols approved by the University Laboratory of Animal Resources (ULAR) of the University of Pennsylvania.

#### Statistical tests

In the ELISA and mouse experiments, all the raw data were log10 transformed and the statistical significance was determined using one-way ANOVA followed by Dunnett’s test. All the p values are shown for each of the groups, all groups were compared to the untreated control group.

#### Statistical analysis

All calculation and statistical analyses of the experimental data were conducted using GraphPad Prism v.10.0.2. Statistical significance between different groups was calculated using the tests indicated in each figure legend. No statistical methods were used to predetermine sample size.

