## Supplementary Information for "Peptides from non-immune proteins target infections through antimicrobial and immunomodulatory properties"

**This PDF file includes:**

Table S1
Figs. S1 to S11

**Table S1. Precursor immune and non-immune proteins of encrypted peptides.** Precursor proteins and their reported functions were obtained from UniProt (https://www.uniprot.org/). Encrypted peptides derived from each of the immune and non-immune proteins are listed.

| **Immune proteins** | **Encrypted peptides** | **Reported function** | **Reference** |
|---|---|---|---|
| Immunoglobulin superfamily member 10 (IGSF10) | Immunosin-1, 2, 3, 4 | It is involved in the control of early migration of neurons expressing gonadotropin-releasing hormone (GNRH neurons) | https://www.uniprot.org/uniprot/Q6WRI0#structure |
| Pro-interleukin-16 (IL16) | Immunosin-5 | It plays a role in the stimulation of a migratory response in CD4+ lymphocytes, monocytes, and eosinophils | https://www.uniprot.org/uniprot/Q14005#structure |
| Leukocyte cell-derived chemotaxin-2 (LECT2) | Immunosin-6 | It has a neutrophil chemotactic activity | https://www.uniprot.org/uniprot/O14960#structure |
| C-C motif chemokine 26 (CCL26) | Immunosin-7 | It is a chemokine receptors binding chemokines, a ligand for C-C chemokine receptor, and it is a chemoattractant for eosinophils and basophils | https://www.uniprot.org/uniprot/Q9Y258#structure |
| T-cell surface glycoprotein CD8 beta chain (CD8B) | Immunosin-8, 9, 10 | It plays role in immunoregulatory interaction between a Lymphoid and a non-Lymphoid cells | https://www.uniprot.org/uniprot/P10966#structure |
| C-C motif chemokine 20 (CCL20) | Immunosin-11 | It acts as a ligand for C-C chemokine receptor CCR6 inducing a strong chemotactic response | https://www.uniprot.org/uniprot/P78556#structure |
| Interleukin-17 receptor E (IL17RE) | Immunosin-12 and 13 | It is a specific functional receptor for IL17C (an autocrine cytokine that regulates innate epithelial immune responses | https://www.uniprot.org/uniprot/Q8NFR9#structure |
| **Non-immune proteins** | | **Reported function** | |
| *Collagen-related* | | | |

| | | | |
|---|---|---|---|
| Collagen alpha-1(XXIV) chain (COOA1) | Collagenin-1 | It is involved in signal transduction, extracellular matrix organization, including assembly of collagen fibrils and collagens biosynthesis. | https://www.uniprot.org/uniprot/Q17RW2#structure |
| Disintegrin and metalloproteinase domain-containing protein 12 (ADAM12) | Collagenin-2 | It is involved in macrophage-derived giant cells (MGC) and osteoclast formation from mononuclear precursors. | https://www.uniprot.org/uniprot/O43184#structure |
| Collagen alpha-1(XVI) chain (COGA1) | Collagenin-3 and 4 | It plays a role in integrin cell-surface interaction, collagen degradation, and collagen biosynthesis; it is also involved in mediating cell attachment and inducing integrin-mediated cellular reactions, such as cell spreading and alterations in cell morphology. | https://www.uniprot.org/uniprot/Q07092#structure |
| Collagen alpha-6(VI) chain (CO6A6) | Collagenin-5, 6 and 7 | Collagen VI acts as a cell-binding protein, and its function is associated with the biosynthesis of collagen. | https://www.uniprot.org/uniprot/A6NMZ7#structure |
| *Neuro-related* | | | |
| Glia-derived nexin (GDN) | Neurin-1 and 2 | It plays a role in the dissolution of fibrin clot and in the inhibition of serine protease with activity toward thrombin, trypsin, and urokinase. It also promotes neurite extension by inhibiting thrombin. | https://www.uniprot.org/uniprot/P07093#structure |
| Neurotrypsin (NETR) | Neurin-3 | It plays a role in neuronal plasticity and the proteolytic action may subserve structural reorganizations associated with learning and memory operations. | https://www.uniprot.org/uniprot/P56730#structure |
| Brain-derived neurotrophic factor (BDNF) | Neurin-4 | BDNF appears to be correlated with Leishmania diseases, HIV and amyloid fiber formation (neurodegenerative diseases). During development, it promotes the survival and differentiation of selected neuronal populations of the peripheral and central nervous systems. It also | https://www.uniprot.org/uniprot/P23560#structure |

| | | | |
|---|---|---|---|
| | | participates in axonal growth, pathfinding and in the modulation of dendritic growth and morphology. It is considered as a major regulator of synaptic transmission and plasticity at adult synapses in many regions of the CNS. The versatility of BDNF is emphasized by its contribution to a range of adaptive neuronal responses including long-term potentiation (LTP), long-term depression (LTD), certain forms of short-term synaptic plasticity, as well as homeostatic regulation of intrinsic neuronal excitability. | |
| Protein NDNF (NDNF) | Neurin-5 and 6 | It is a secretory protein that plays a role in various cellular processes. It also acts as a chemorepellent acting on gonadotropin-releasing hormone (GnRH) expressing neurons regulating their migration to the hypothalamus. Moreover, this protein promotes neuron migration, growth and survival as well as neurite outgrowth and is involved in the development of the olfactory system. It may also act through the regulation of growth factors activity and downstream signaling. It has found to regulate extracellular matrix assembly and cell adhesiveness (By similarity) and to promote endothelial cell survival, vessel formation and plays an important role in the process of revascularization through NOS3-dependent mechanisms. | https://www.uniprot.org/uniprot/Q8TB73#structure |
| *Eye-related* | | | |
| BMP/retinoic acid-inducible neural-specific | Eyesin-1, 2, 3 and 4 | It inhibits neuronal cell proliferation by negative regulation of the cell cycle transition, and it promotes pituitary gonadotrope cell proliferation, migration and invasion, when overexpressed. It | https://www.uniprot.org/uniprot/Q76B58#structure |

| | | | |
|---|---|---|---|
| protein 3 (BRNP3) | | may play a role in cell pituitary tumor development. | |
| BMP/retinoic acid-inducible neural-specific protein 2 (BRNP2) | Eyesin-5 and 6 | It inhibits neuronal cell proliferation by negative regulation of the cell cycle transition. | https://www.uniprot.org/uniprot/Q9C0B6#structure |
| Interphotoreceptor matrix proteoglycan 1 (IMPG1) | Eyesin-7 and 8 | It is defined as chondroitin sulfate-, heparin- and hyaluronan-binding protein (By similarity). It may serve to form a basic macromolecular scaffold comprising the insoluble interphotoreceptor matrix. | https://www.uniprot.org/uniprot/Q17R60#structure |
| *Odontogenesis-related* | | | |
| Odontogenesis associated phosphoprotein (ODAPH) | Toothin-1, 2 and 3 | It may promote nucleation of hydroxyapatite. | https://www.uniprot.org/uniprot/Q17RF5#structure |
| *Leucine-zipper-related* | | | |
| Leucine-rich repeat-containing protein 17 (LRRC17) | Zipperin-1 and 2 | It is involved in bone homeostasis. It also acts as a negative regulator of RANKL-induced osteoclast precursor differentiation from bone marrow precursors (By similarity). | https://www.uniprot.org/uniprot/Q8N6Y2#structure |

*Supplementary Figures*

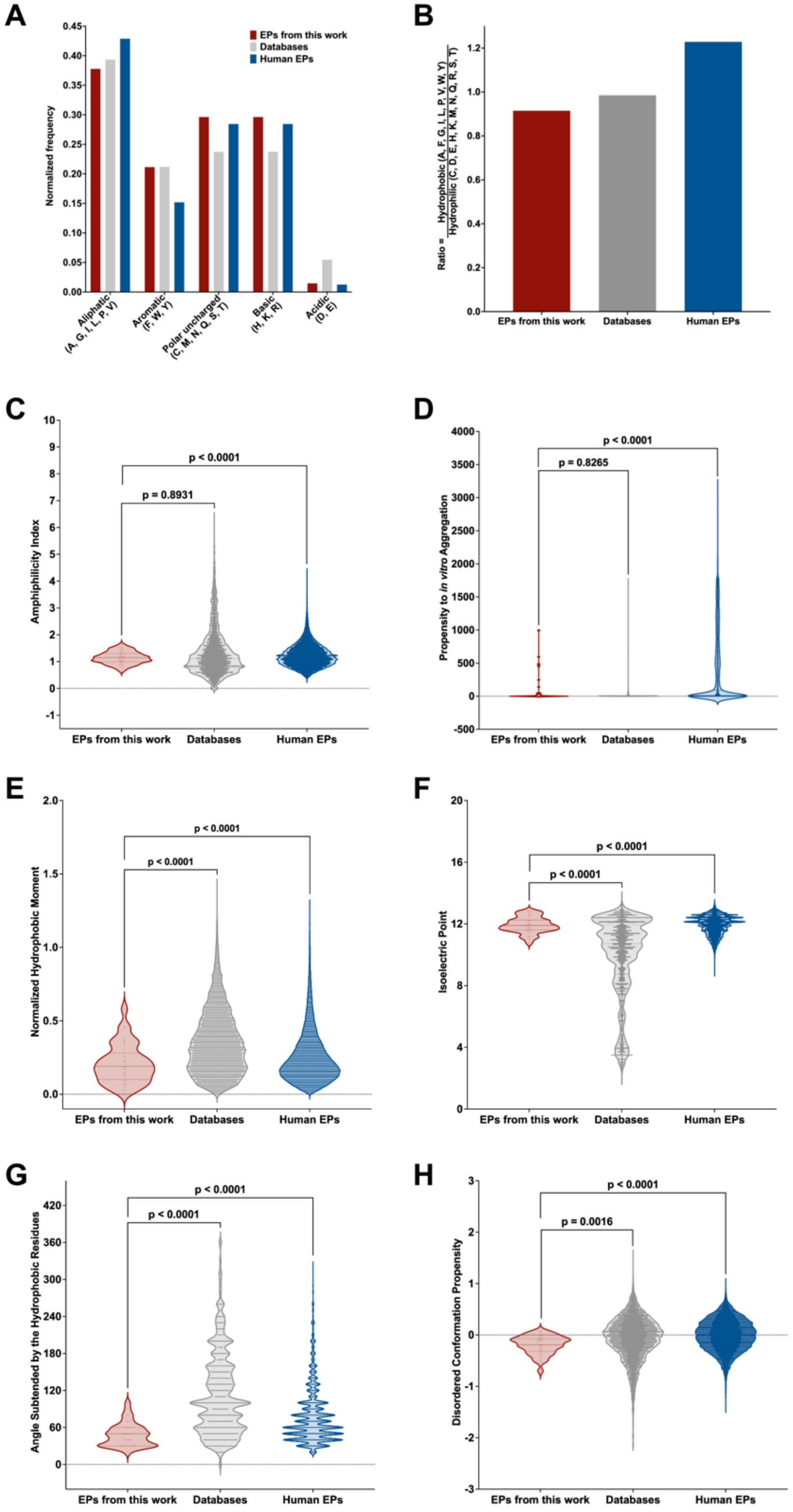


**Fig. S1. Frequency distribution by amino acid type and physicochemical features of EPs compared to known AMPs and other EPs from the human proteome. (A)** Normalized frequency of amino acid residue types of the EPs from the six biogeographical areas compared to AMPs from databases (DBAASP, APD3, and DRAMP) and all EPs from the human proteome. **(B)** Ratio between Hydrophilic and Hydrophobic amino acid residues for each one of the three different classes of peptides (EPs from this work, AMPs, and EPs from the human proteome). The following physicochemical features were estimated using the Database of Antimicrobial Activity and Structure of Peptides (DBAASP) server: **(C)** amphiphilicity index, **(D)** propensity to in vitro aggregation, **(E)** normalized hydrophobic moment, **(F)** isoelectric point, **(G)** angle subtended by the hydrophobic residues, and **(H)** disordered conformation propensity. Those physicochemical features and net charge (**Fig. 1B**) and normalized hydrophobic moment (**Fig. 1C**) are the most relevant responsible for directly influence the antimicrobial activity and toxicity of peptides.

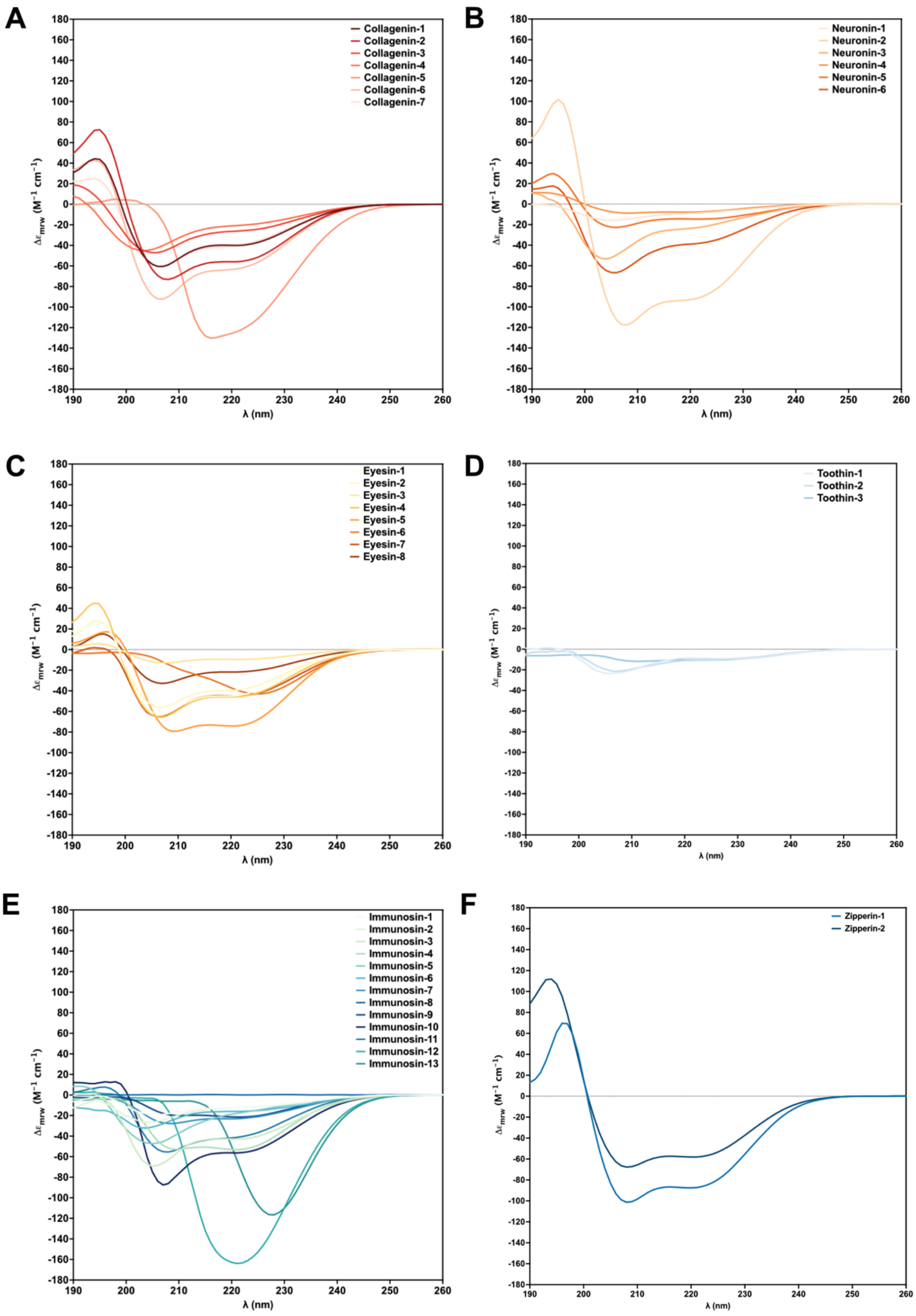


**Fig. S2. Secondary structure of EPs.** Circular Dichroism spectra of **(A)** collagenins, **(B)** neuronins, **(C)** eyesins, **(D)** toothins, **(E)** immunosins, and **(F)** zipperins. The experiments were performed in trifluoroethanol (TFE) and water (3:2, v:v) mixture. Experimental conditions are described in the ***Method details*** section.

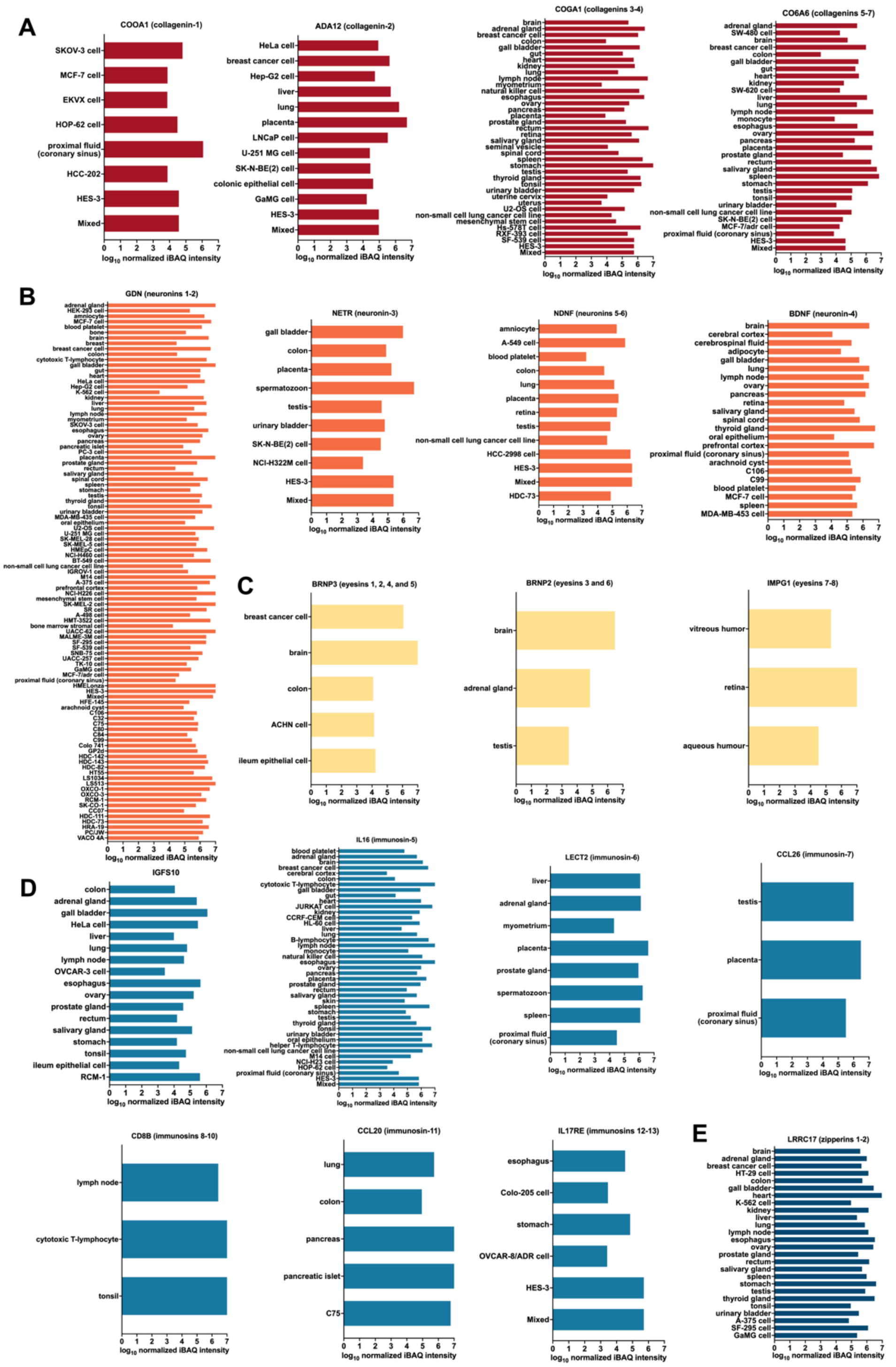


**Fig. S3. Expression levels of the parent proteins of EPs from the human proteome.** Normalized expression level values expressed in $\log_{10}$ intensity based absolute quantification (iBAQ), a metric for parent protein expression and abundance of **(A)** collagenins, **(B)** neuronins, **(C)** eyesins, **(D)** immunosins, and **(E)** zipperins. The parent protein of toothins (Odontogenesis associated phosphoprotein) does not have expression levels reported in the ProteomicsDB server (https://www.proteomicsdb.org/).

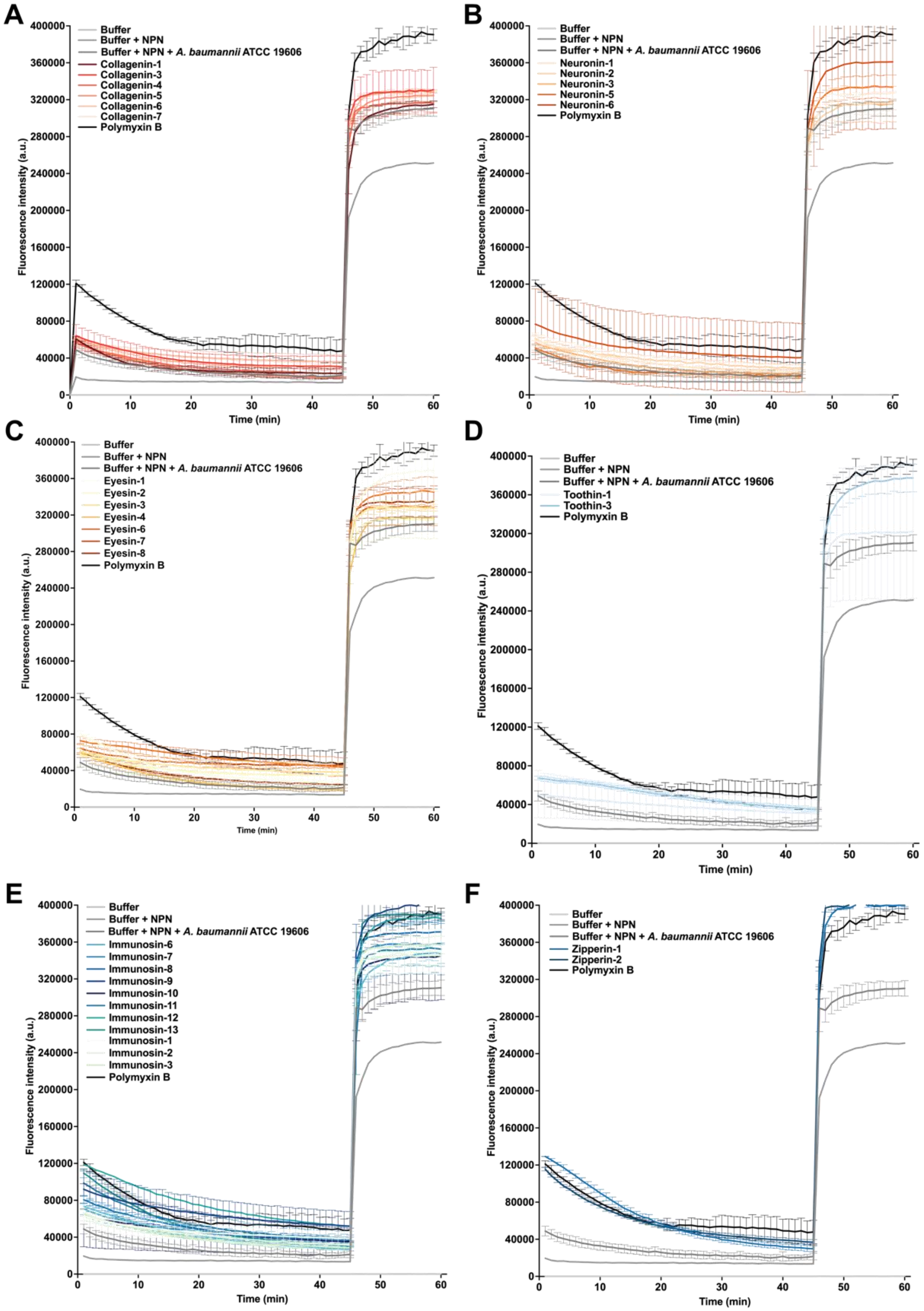


**Fig. S4. Outer membrane permeabilization experiments with EPs.** Permeabilization of the outer membrane using the probe 1-(N-phenylamino)naphthalene (NPN) on the Gram-negative strain *A. baumannii* ATCC 19606 of **(A)** collagenins, **(B)** neuronins, **(C)** eyesins, **(D)** toothins, **(E)** immunosins, and **(F)** zipperins.

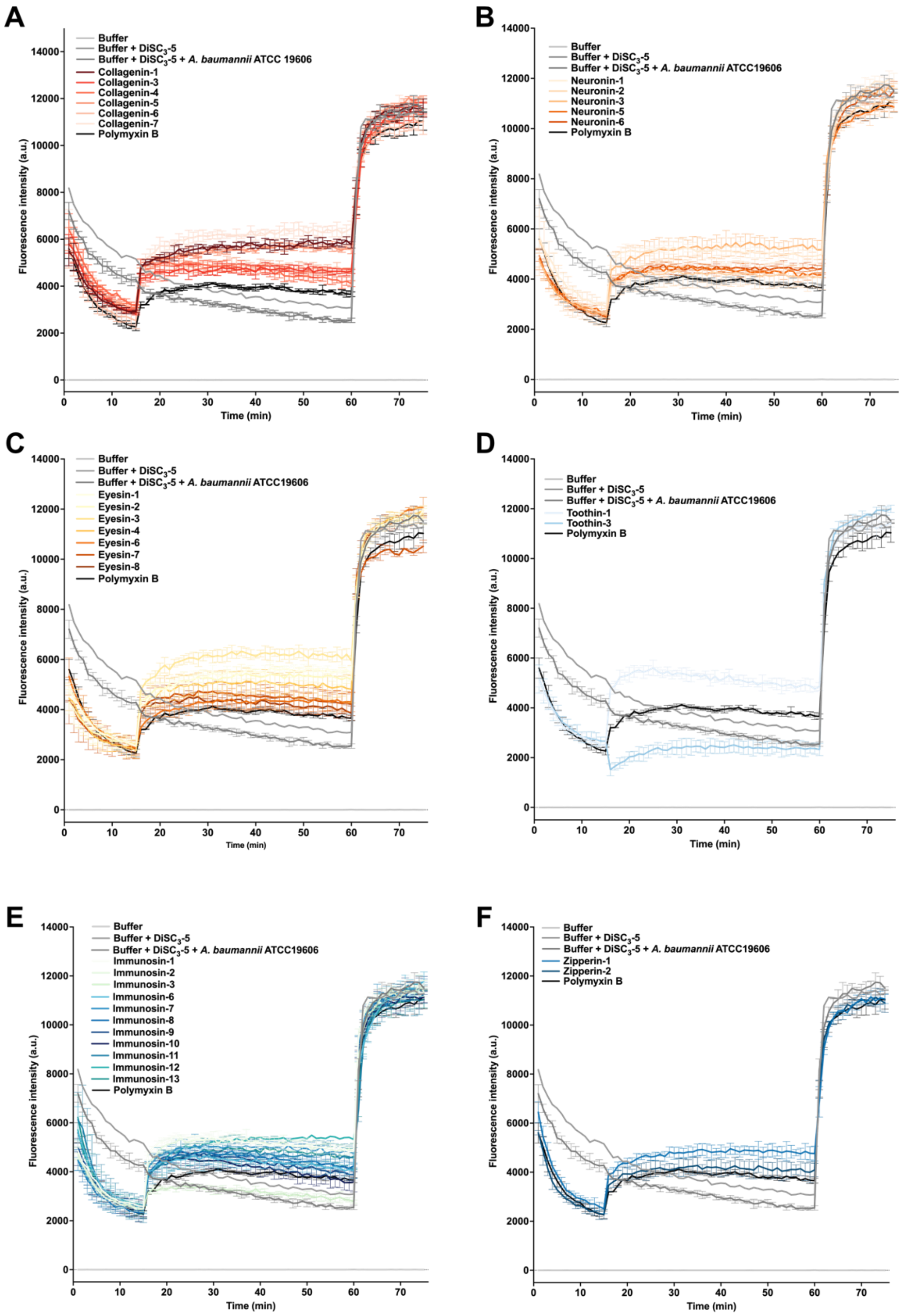


**Fig. S5. Cytoplasmic membrane depolarization triggered by EPs on *A. baumannii* cell membranes.** Depolarization assays with the hydrophobic probe 3,3′-dipropylthiadicarbocyanine iodide (DiSC$_3$-5) on the cytoplasmic membrane of *A. baumannii* ATCC 19606 cells of **(A)** collagenins, **(B)** neuronins, **(C)** eyesins, **(D)** toothins, **(E)** immunosins, and **(F)** zipperins.

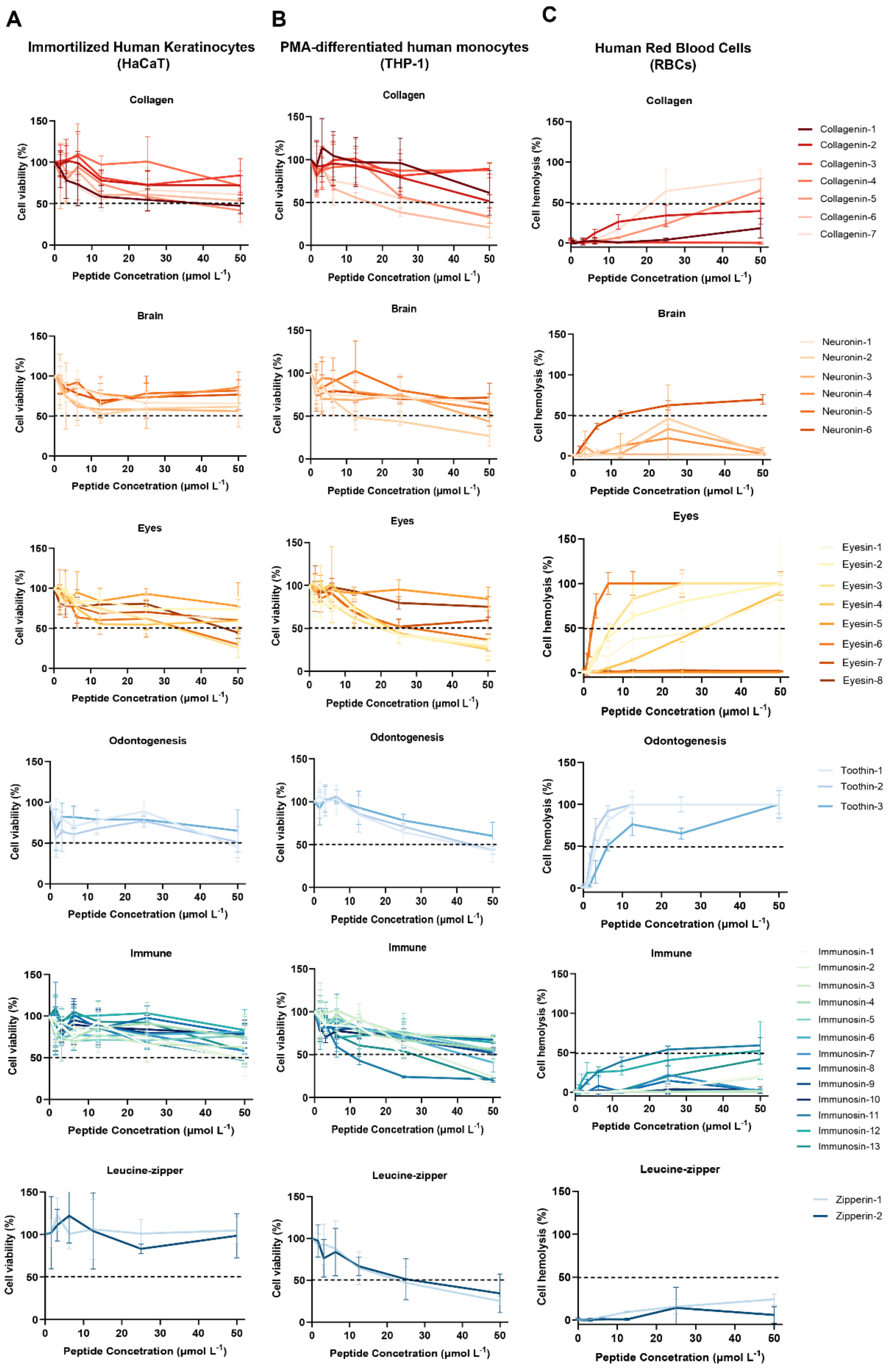


**Fig. S6.** Cytotoxicity and hemolytic effects of the selected EPs. Cytotoxic effects of increasing concentrations (0, 3.12, 6.25, 12.5, 25, and 50 μmol $L^{-1}$) of each EP on **(A)** immortalized human keratinocytes (HaCaT) and **(B)** PMA-differentiated human monocytes (THP-1) after 24 h of treatment. **(C)** Analysis of hemolytic effects of each EP on human red blood cells (RBCs) after 1 h of incubation at 37 °C. Assays have been performed in triplicate and data represent the average of the cell viability obtained by comparing each sample with the untreated cells (control).

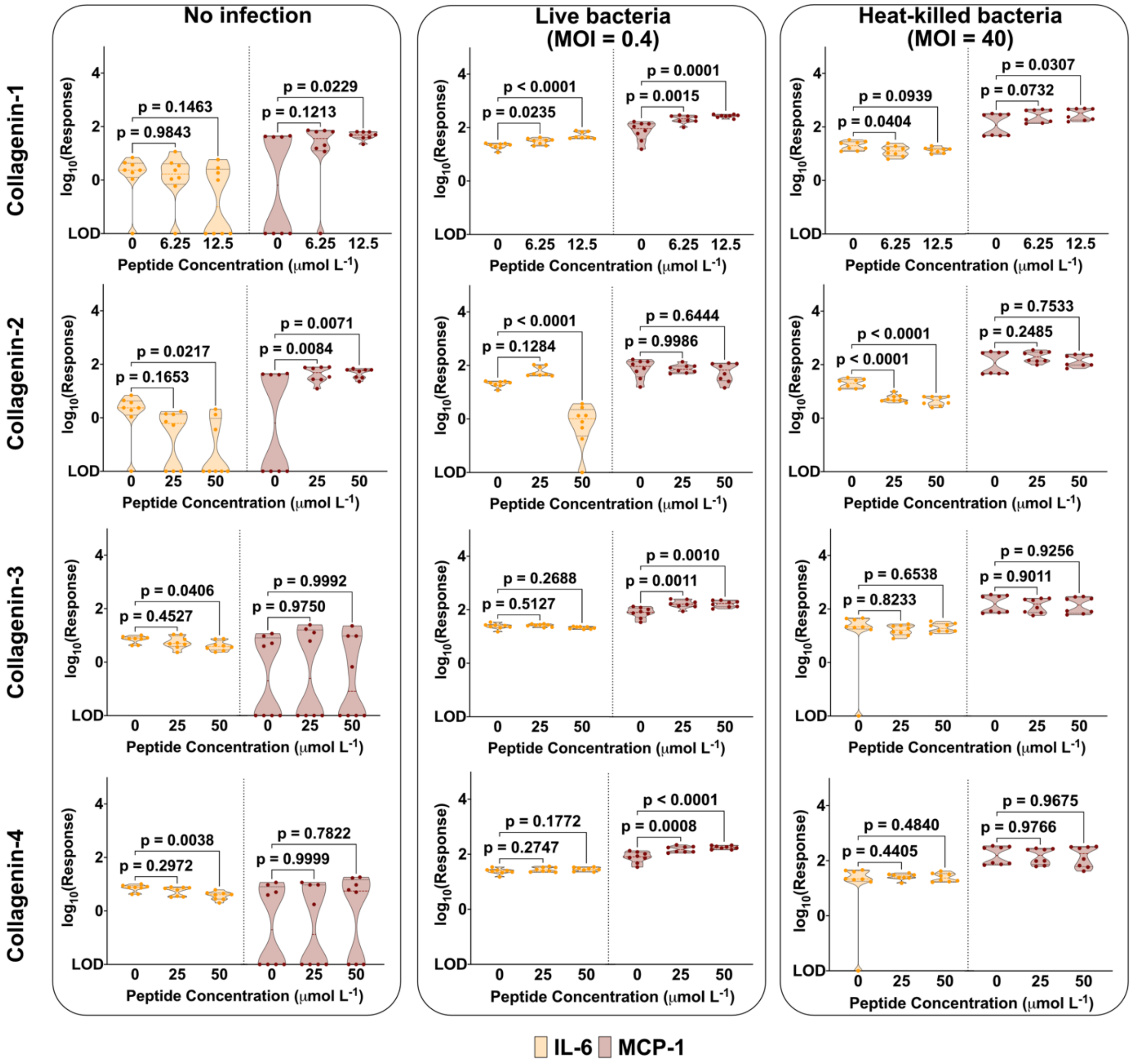


**Fig. S7. Immunomodulatory properties of collagen-derived EPs on keratinocytes.** Effects of collagenin-1, 2, 3, and 4 on the release of IL-6 and MCP-1 by HaCaT cells, both uninfected and infected with live or heat-killed *A. baumannii* ATCC 19606. To determine statistical significance, we used one-way ANOVA followed by Dunnett's test, and the respective p-values are presented for each group. All groups were compared to the untreated control, and the violin plots display median and upper and lower quartiles. Values beyond the lower limit of detection of the ELISA assay were noted as LOD.

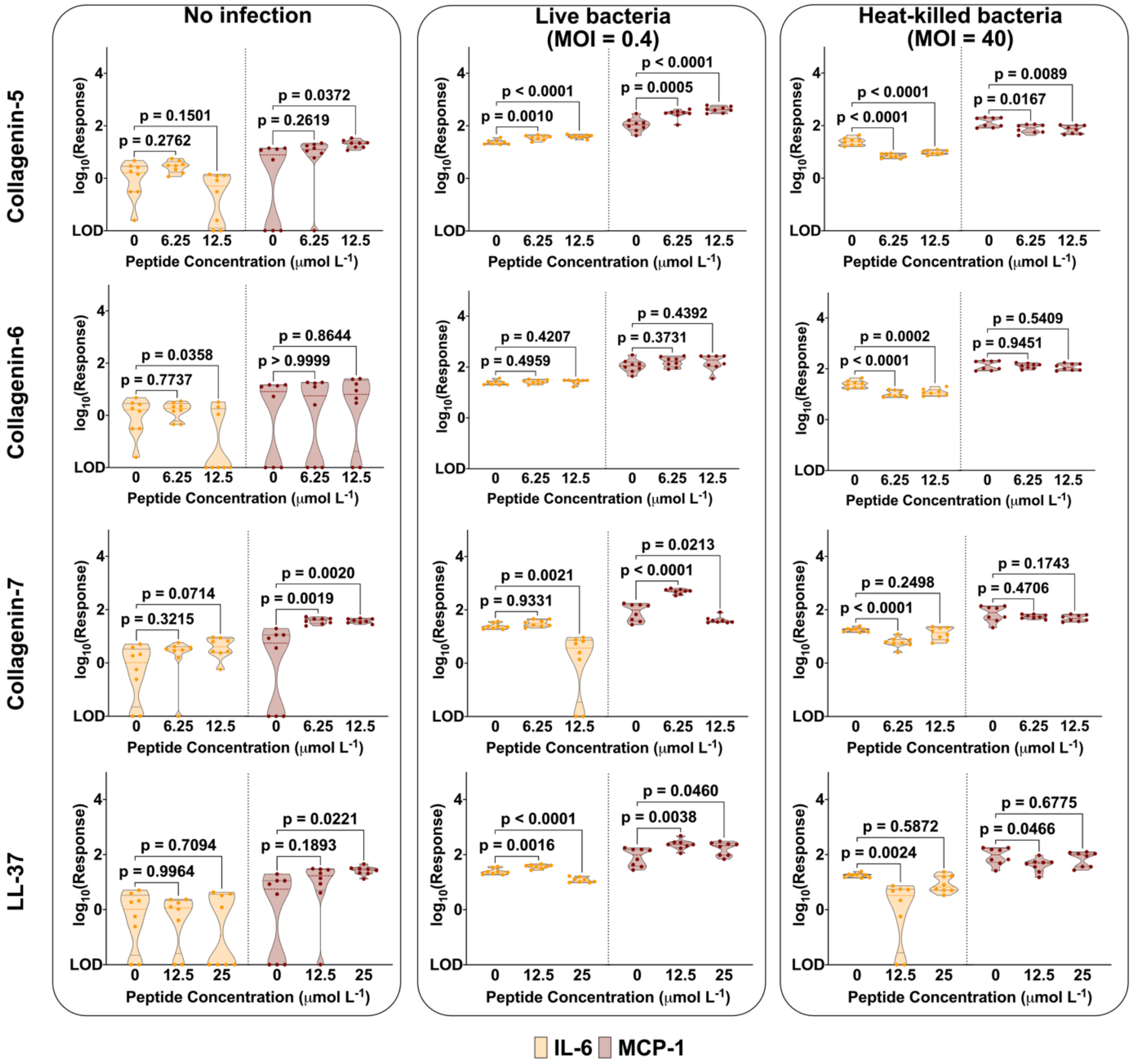


**Fig. S8. Immunomodulatory properties of collagen derived EPs on keratinocytes.** Effects of collagenin- 5, 6, 7, 8, and LL-37 on the release of IL-6 and MCP-1 by HaCaT cells, both uninfected and infected with either live or heat-killed *A. baumannii* ATCC 19606. Statistical significance was determined as detailed in the legend of **Fig. S7**. Values beyond the lower limit of detection of the ELISA assay were noted as LOD.

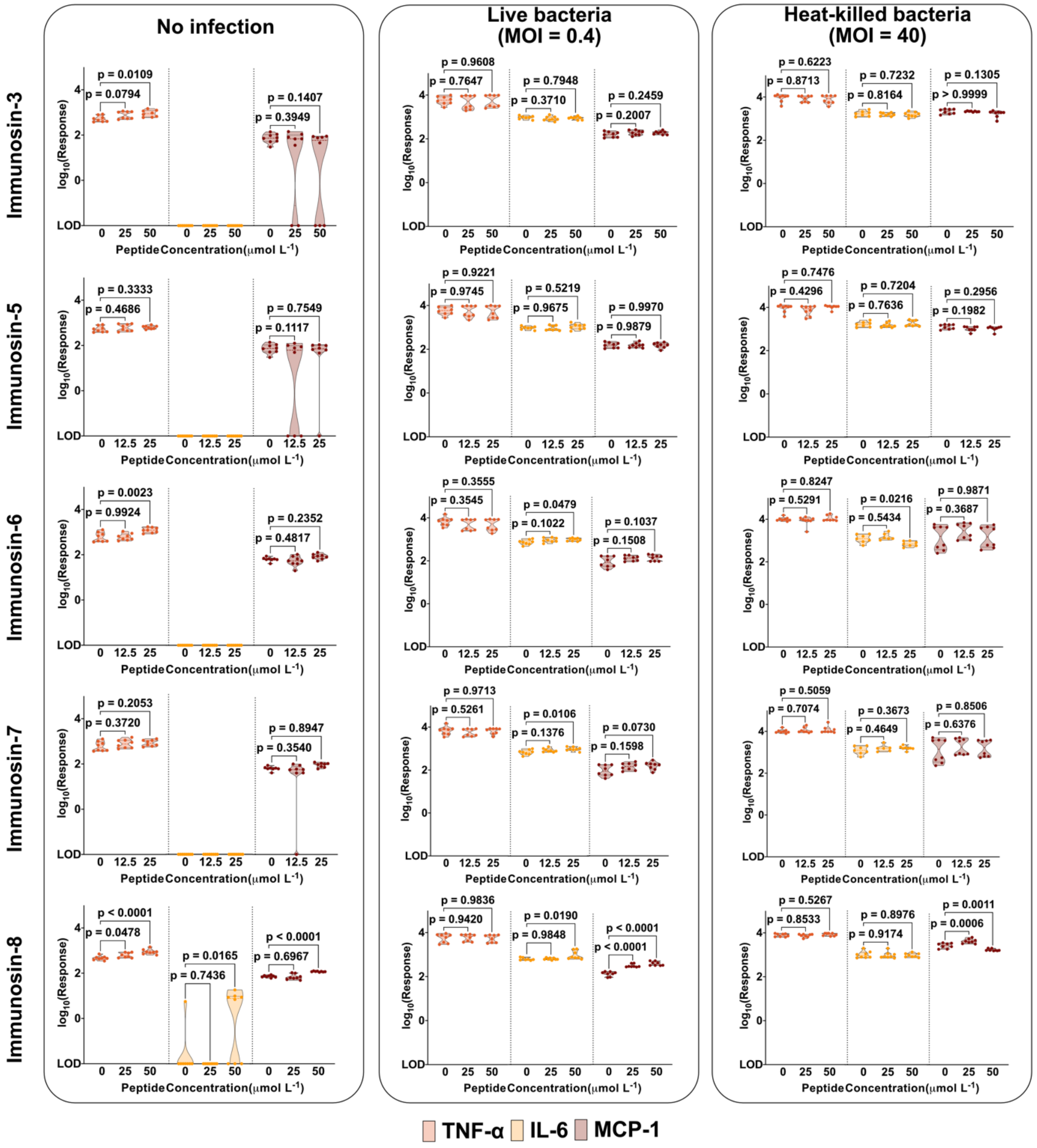


**Fig. S9. Immunomodulatory properties of immune system and leucine zipper-derived EPs on PMA-differentiated human monocytes.** Effects of immunosin-3, 5, 6, 7, and 8 on the release of TNF-α, IL-6 and MCP-1 by phorbol 12-myristate-13-acetate (PMA)-differentiated THP-1, both uninfected and infected with either live or heat-killed *A. baumannii* ATCC 19606. Statistical significance was determined as detailed in the legend of **Fig. S7**. Values beyond the lower limit of detection of the ELISA assay were noted as LOD.

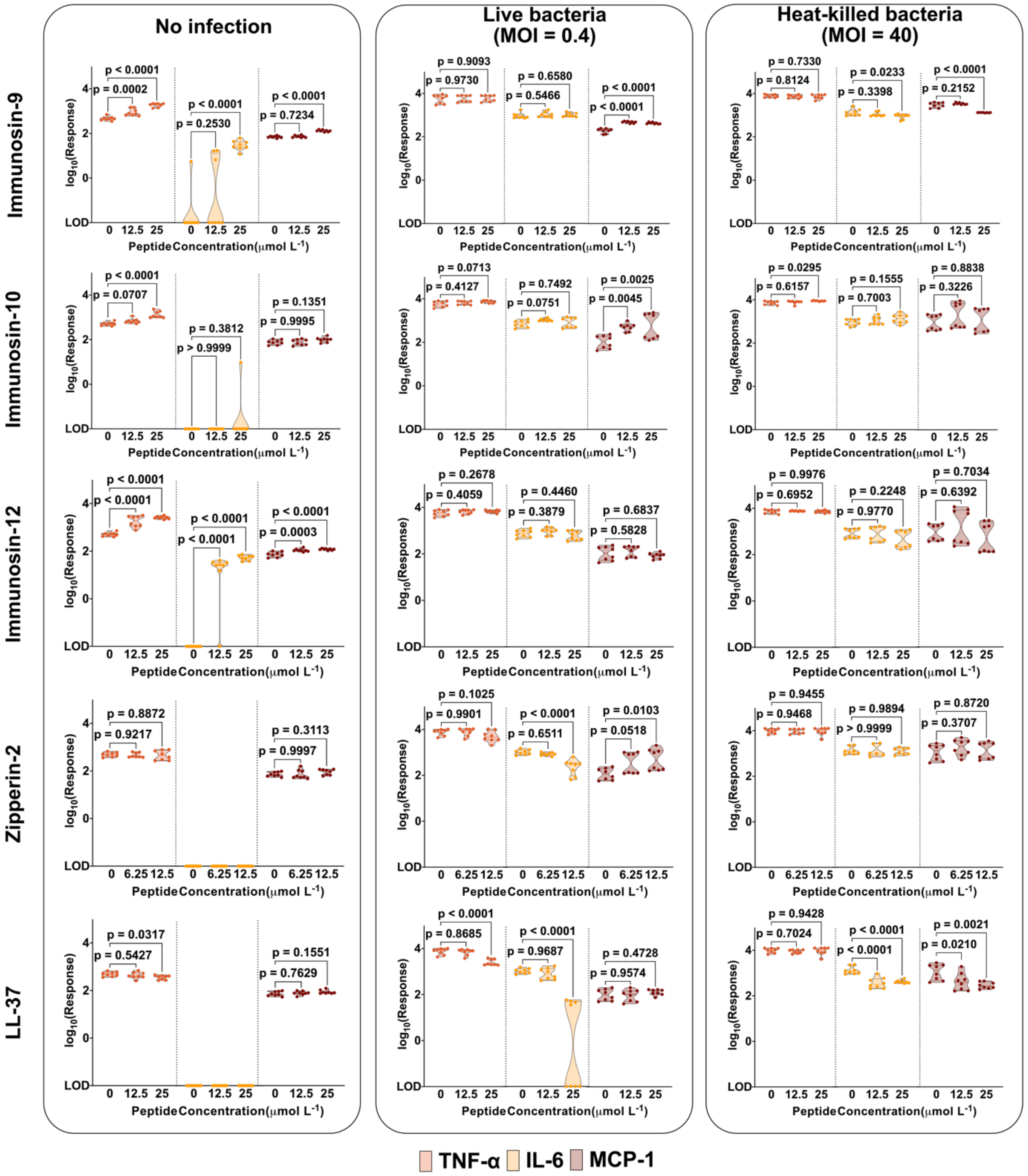


**Fig. S10. Immunomodulatory properties of immune system- and leucine zipper-derived EPs on PMA-differentiated human monocytes.** Effects of immunosin-9, 10, 11, zipperin-2, and LL-37 on the release of tnf-α, IL-6 and MCP-1 by phorbol 12-myristate-13-acetate (PMA)-differentiated THP-1, both uninfected and infected with either live or heat-killed *A. baumannii* ATCC 19606. Statistical significance was determined as detailed in the legend of **Fig. S7**. Values beyond the lower limit of detection of the ELISA assay were noted as LOD.

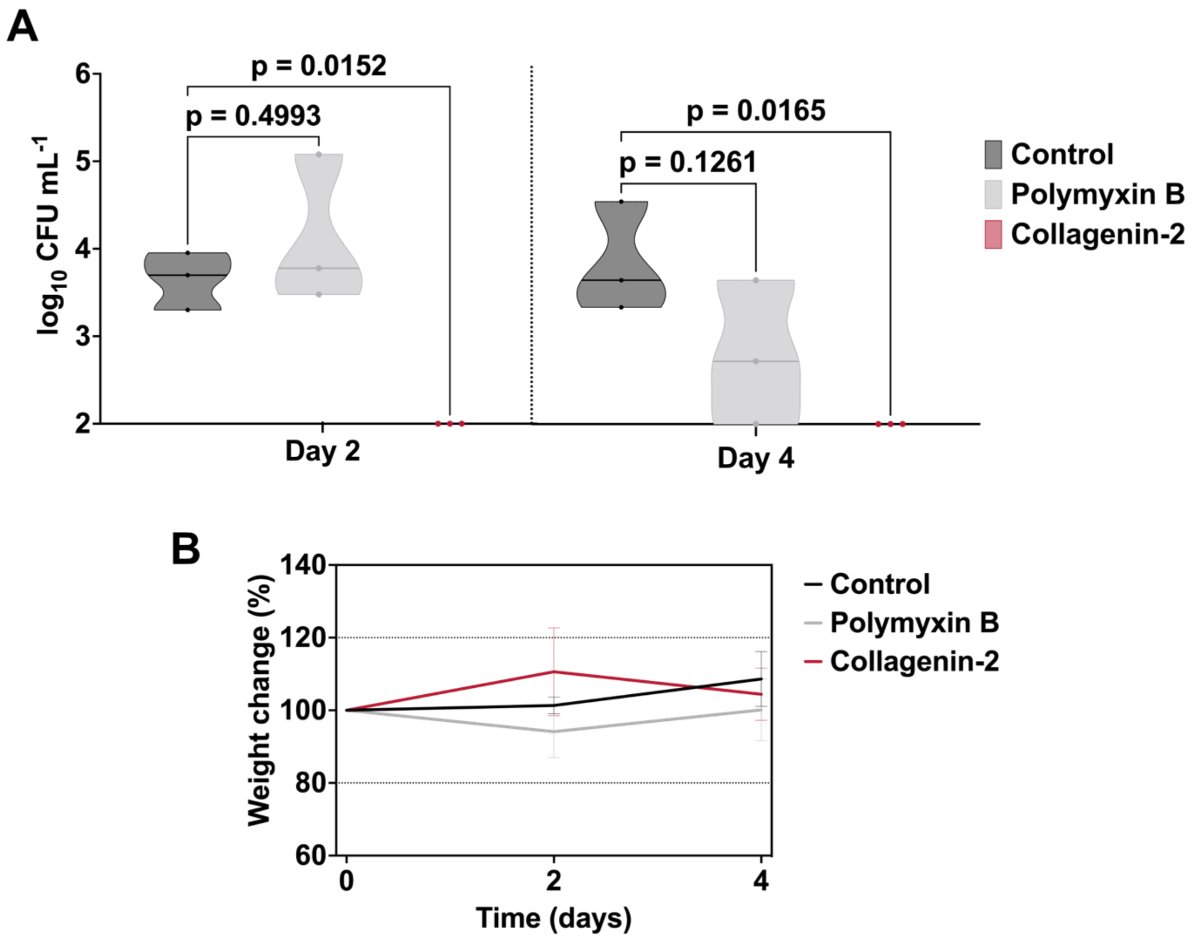


**Fig. S11. Anti-infective activity of collagenin-2 in skin abscess infection mouse model.** In this model, the back of the mice was shaved, wounded, and infected with *A. baumannii* cells. One hour after the infection a single dose of collagenin-2 at 50 µmol $L^{-1}$ was applied to the wounded epidermis. **(A)** Two- and four-days post-infection mice were euthanized, and their skin tissue was collected for bacterial counting. **(B)** During the experiment, the weight was monitored to rule out the toxicity of the EPs. We considered 20% weight change as the threshold for lack of toxicity.
